# Studying the fate of tumor extracellular vesicles at high spatio-temporal resolution using the zebrafish embryo

**DOI:** 10.1101/380238

**Authors:** Vincent Hyenne, Shima Ghoroghi, Mayeul Collot, Sébastien Harlepp, Jack Bauer, Luc Mercier, Ignacio Busnelli, Olivier Lefebvre, Nina Fekonja, Pedro Machado, Joanna Bons, François Delalande, Ana Isabel Amor, Susana Garcia Silva, Frederik J. Verweij, Guillaume Van Niel, Yannick Schwab, Héctor Peinado, Christine Carapito, Andrey S. Klymchenko, Jacky G. Goetz

## Abstract

Tumor extracellular vesicles (tumor EVs) mediate the communication between tumor and stromal cells mostly to the benefit of tumor progression. Notably, tumor EVs have been reported to travel in the blood circulation, reach specific distant organs and locally modify the microenvironment. However, visualizing these events *in vivo* still faces major hurdles. Here, we show a new method for tracking individual circulating tumor EVs in a living organism: we combine novel, bright and specific fluorescent membrane probes, MemBright, with the transparent zebrafish embryo as an animal model. We provide the first description of tumor EVs’ hemodynamic behavior and document their arrest before internalization. Using transgenic lines, we show that circulating tumor EVs are uptaken by endothelial cells and blood patrolling macrophages, but not by leukocytes, and subsequently stored in acidic degradative compartments. Finally, we prove that the MemBright can be used to follow naturally released tumor EVs *in vivo*. Overall, our study demonstrates the usefulness and prospects of zebrafish embryo to track tumor EVs *in vivo*.

**Highlights:**

- MemBright, a new family of membrane probes, allows for bright and specific staining of EVs
- Zebrafish melanoma EVs are very similar to human and mouse melanoma EVs in morphology and protein content
- The zebrafish embryo is an adapted model to precisely track tumor EVs dynamics and fate in a living organism from light to electron microscopy
- Circulating tumor EVs are rapidly uptaken by endothelial cells and patrolling macrophages
- Correlated light and electron microscopy can be used in zebrafish to identify cells and compartments uptaking tumor EVs

**Blurb:** Dispersion of tumor extracellular vesicles (EVs) throughout the body promotes tumor progression. However the behavior of tumor EVs in body fluids remains mysterious due to their small size and the absence of adapted animal model. Here we show that the zebrafish embryo can be used to track circulating tumor EVs *in vivo* and provide the first high-resolution description of their dissemination and uptake.

## Introduction

Over the past two decades, extracellular vesicles (EVs) have emerged as novel mediators of cell-cell communication in a wide range of living organisms and biological contexts, due to their capacity to carry functional molecules coupled to their ability to travel in biological fluids (Raposo and Stoorvogel, 2013). EVs are heterogeneous in content and origin, as they can either arise from plasma membrane budding (then called microvesicles) or originate from a late endosomal compartment, the multi-vesicular body (MVB) (then called exosomes) (Kowal et al., 2014; van Niel et al., 2018). EVs are known to be important in tumor progression and metastasis, where the complex tumoral microenvironment requires a permanent cross-communication between cells (Becker et al., 2016; Hyenne et al., 2017). EVs secreted by tumor cells are enriched in pro-tumoral and pro-metastatic factors (proteins, mRNAs, miRNAs and other non-coding RNAs) and can modify the phenotype of both tumor and stromal cells that have uptaken them, mostly to the benefit of tumor growth and metastasis formation (Becker et al., 2016; Hyenne et al., 2017). For instance, tumor EVs were shown to transfer oncogenic traits from more aggressive to less aggressive tumor cells (Al-Nedawi et al., 2008; Chen et al., 2014; Di Vizio et al., 2009). Importantly, tumor EVs can transform macrophages or fibroblasts into tumor associated macrophages or fibroblasts, thereby promoting tumor growth and invasion (Chow et al., 2014; Gu et al., 2012; Paggetti et al., 2015). This EV-mediated communication can occur locally within the primary tumor or at distance in physically far-off organs where it fosters metastatic progression (Peinado et al., 2017). Remarkably, several studies demonstrated that repeated injection of EVs isolated from metastatic cells into the mouse blood circulation induces the formation of a pre-metastatic niche, even in the absence of tumor cells (Costa-Silva et al., 2015; Grange et al., 2011; Hoshino et al., 2015; Liu et al., 2016; Peinado et al., 2012). The ability of circulating tumor EVs to alter the microenvironment of a given distant organ is particularly relevant with regards to i) the increased amounts of tumor EVs present in the blood circulation of patients with cancer (Baran et al., 2010; Galindo-Hernandez et al., 2013; Logozzi et al., 2009) and ii) the fact that elevated levels of EV proteins have been associated with poor prognosis in metastatic melanoma patients (Peinado et al., 2012). Therefore, it is crucial to precisely understand the mechanisms governing tumor EV dispersion and uptake in the blood circulation.

However, local or distant dissemination of tumor EVs has only been sparsely characterized in living organisms (Hoshino et al., 2015; Lai et al., 2015; Pucci et al., 2016). In particular, how EVs circulate in the blood flow, how they cross the endothelial barrier or how specifically they are uptaken by stromal cells during the priming of pre-metastatic niches remains poorly understood due to the lack of adapted and synergistic animal model and imaging approaches. EVs are tens of nanometers sized objects and are thus difficult to track *in vivo*. Moreover, the most studied animal model for tumor growth, the mouse, is not fully suited for real time and *in vivo* EV tracking. In living mice, EVs can either be followed after bulk injections at the level of the whole organism (Lai et al., 2014; Takahashi et al., 2013) or, more precisely through efficient yet relatively complex intravital imaging procedures (Lai et al., 2015; Van Der Vos et al., 2016; Zomer et al., 2015). Such approaches have not yet been able to describe the behavior of tumor EVs in the blood circulation, while this is a step of utmost importance as circulating EVs establish the dialog with the distant microenvironment. An ideal animal model suited to accurately study and understand the behavior of tumor EVs *in vivo* would allow their tracking after release in the circulation and their uptake and, in the same time, be amenable for modeling tumor and metastasis progression.

The zebrafish embryo combines these different characteristics. Indeed, zebrafish has recently emerged as a potent model in cancer biology, as it can be used to study the main steps of tumor progression, such as primary tumor growth, intravasation, extravasation and metastasis *in vivo* (White et al., 2013). The molecular pathways driving cancer progression and the anatomo”pathological features of tumorigenesis are essentially conserved between human and fish (White et al., 2013). In addition, the zebrafish embryo is transparent, possesses a stereotyped vasculature, a maturating immune system and is therefore perfectly suited for intravital imaging with high spatial and temporal resolution. For these reasons, the zebrafish embryo appears as an adequate model to study tumor EVs *in vitro*. In order to track EVs, different fluorescent labeling methods have been described and debated (Hyenne et al., 2017). Genetic encoding of fusion proteins between fluorophores and proteins/peptides enriched in EVs presents the advantage of being specific. However, EVs are now known to be highly heterogeneous in content (Kowal et al., 2016) and such approaches are therefore often restricted to subpopulations of vesicles. In addition, they are not applicable to EVs derived, for example, from cancer patients’ samples. As an alternative, membrane binding fluorescent dyes can be used to rapidly and brightly label any EV populations regardless of their origin. However, they are not specifically designed to label EVs and their specificity has been questioned (Lai et al., 2014; Takov et al., 2017).

Here, we use two recently developed cyanine-based membrane probes, called MemBright, bearing alkyl chains and zwitterionic groups MemBright-Cy3 and MemBright-Cy5, for efficient staining of EVs (Collot et al., 2018), submitted). These probes exhibit drastic improvement in terms of brightness and specificity of EVs staining as compared to commonly used PKH-26. Besides, we show that zebrafish melanoma EVs are similar in morphology and protein content to human melanoma EVs and demonstrate how their fate can be tracked in the zebrafish embryo. Upon injection in the blood circulation, we successfully tracked individual flowing EVs using high-speed confocal imaging. We provide the first description of EVs’ dynamics in the blood circulation. We subsequently examined the transit routes and arrest sites of tumor EVs and identified endothelial cells and patrolling macrophages as main recipient cells. Importantly, these cells have been similarly identified in a parallel study describing endogenous EVs dispersion in the zebrafish embryo (Verweij et al., co-submitted). We found that patrolling macrophages internalize tumor EVs through at least two distinct endocytic mechanisms. In addition, after internalization, EVs are stored in acidic compartments. Using correlated light and electron microscopy, we were able to precisely identify the cells uptaking EVs and finely describe their morphology as well as the storage/degradative compartments at the electron microscopy level. Finally, we demonstrate that cells stained with MemBright can naturally release labeled exosomes, which can be further tracked *in vitro* and *in vivo* in the zebrafish embryo.

## Results

### Zebrafish melanoma EVs are similar to human and mouse melanoma EVs

To study tumor EVs in zebrafish, we first characterized EVs released by a melanoma cell line (Zmel1) derived from a transgenic mitfa-BRAF(V600E);p53(-/-) zebrafish line (Heilmann et al., 2015) (Fig.1A). EVs were isolated from cell culture supernatant following an established protocol of differential centrifugation (Théry et al., 2006) and EVs present in the 100.000g pellet were characterized by NTA (Nanosight, Malvern) and electron microscopy. We found that Zmel1 EVs have an average diameter of 150nm in solution and 90nm after chemical fixation (Fig.1B-C). Subsequently, we characterized the protein content of these EVs by mass spectrometry and identified 544 proteins present in Zmel1 EVs (Table 1). This list includes several proteins typically found in extracellular vesicles, such as Alix, CHMP4C, TSG101 CD9, CD81, Flotillin 1, syntenin 2, integrins α5 and β1 and others (of note, CD63 was absent from Zmel1 EVs) (Fig. 1D and Table 1). We then wondered whether the content of zebrafish melanoma EVs was comparable to the ones of human or mice melanoma EVs. To address this question, we compared proteins present in Zmel1 EVs with proteins identified in the EVs isolated from 6 human (451-LU, SK-Mel28, SK-Mel147, SK-Mel103, WM35, WM164) (Table 2) and 3 mouse (B16-F0, B16-F1, B16-F10) (Table 3) melanoma cell lines. Protein content comparison revealed that among the proteins with orthologs in all three species, respectively 70% and 46% of Zmel1 proteins were also identified in human or mice melanoma EVs (Fig. 1E). By contrast, less than 10% of Zmel1 EV proteins were found in EVs from other human skin cell lines (protein lists obtained from EVpedia; Fig. 1E). We identified a core list of 84 proteins found in melanoma EVs from either zebrafish, mice or human (Table 4). Altogether, these data demonstrate that Zmel1 EVs derived from an established zebrafish melanoma cell line are highly similar to mammalian melanoma EVs and therefore constitute a good model to study human melanoma EVs.

**Figure 1:**
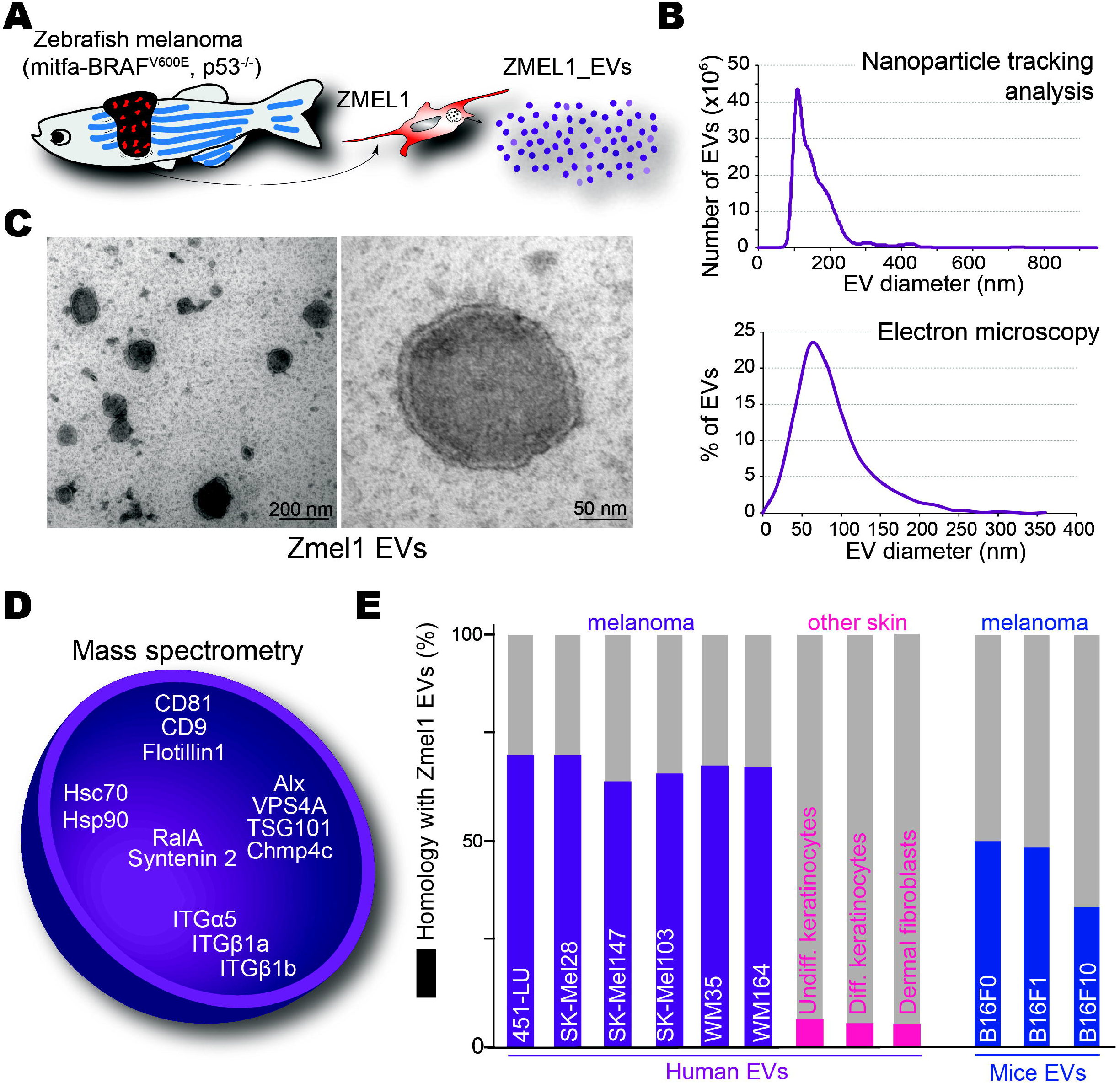
EVs secreted by Zmel1 zebrafish melanoma cells are similar to mouse and human melanoma EVs. **(A)** Zebrafish melanoma EVs were isolated from Zmel1 cells (described by Heilmann and colleagues 2015) by differential centrifugation. **(B)** Representative histogram of a Nanosight nanoparticle tracking analysis of EVs produced by Zmel1 cells showing the number of EVs (y axis) versus their diameter (nm, x axis). **(C)** Representative electron microscopy images of Zmel1 EVs and a histogram showing percentage of total EVs (y axis) versus their diameter (nm, x axis). **(D)** Illustration of some of the classical EV proteins present among the 544 proteins identified in ZMel1 EVs by mass spectrometry (full list is available in Table 1). **(E)** Histogram showing the percentage of Zmel1 EVs proteins common with EV proteins from various human or mouse cell lines (using human orthologs).

In addition, we compared proteins present in Zmel1 EVs with proteins present in two other types of zebrafish EVs identified in a parallel study (Verweij *et al*. co-submitted). First, we found that 26% of Zmel1 EVs proteins are also present in EVs from AB9 fibroblastic cell line (Table 5). Then, we compared Zmel1 EVs with CD63-positive EVs secreted by a zebrafish embryonic epithelium, the yolk syncytial layer (YSL), and isolated from zebrafish embryos (Verweij *et al*. co-submitted). Interestingly, we found a relatively low similarity between these two types of zebrafish EVs (2,8% of Zmel1 EV proteins are present in YSL CD63+ EVs; 14,7% of YSL CD63+ EV proteins are present in Zmel1 EVs) (common proteins listed in Table 6). These results show that the content of EVs secreted by Zmel1 cells differs from CD63-positive EVs secreted by YSL during embryonic development, illustrating the cell type specificity of EV cargo enrichment. However, the mechanism of biogenesis of these two EV types could be partially similar, as 6 of the 15 proteins common to Zmel1 EVs and YSL EVs have been shown to affect, positively or negatively, exosome secretion in mammalian cells: CHMP4C, Alix, Syntenin 2, Flotillin 1, Rab2 and CDC42 (Baietti et al., 2012; Colombo et al., 2013; Ekström et al., 2014; Kajimoto et al., 2017; Okabayashi and Kimura, 2010; Ostrowski et al., 2010; Schulz et al., 2015).

### The MemBright dye specifically and brightly labels tumor EVs

In order to fluorescently label Zmel1 EVs and follow them *in vivo*, we used novel membrane probes, MemBright, that were recently developed (Collot et al., 2018). They differ significantly from existing commercial dyes because they bear two amphiphilic groups composed of zwitterions and alkyl chains, which insert the dye into the membrane bilayer (Fig. 2A). Moreover, MemBright is available in several colors, which therefore enables multi-color approach in EV imaging (See latter, supplementary Figure 5). To assess the value of MemBright in EV labeling, we first globally compared the MemBright-labeled EVs to identical EVs labeled with PKH-26, a commercially available and widely used dye for EV labeling (Christianson et al., 2013; Hoshino et al., 2015; Imai et al., 2015; Tominaga et al., 2015). Zmel1 EVs were incubated with MemBright-Cy3 (at 0,2μM) or with PKH-26 (at 2μM, according to manufacturer’s instructions), washed in a large volume of PBS and isolated by ultracentrifugation. Using fluorescence spectroscopy, we observed that PKH-labeled EVs display a broad absorption spectrum, with a blue shifted peak typically indicating the presence of H-aggregation (Fig. 2B) (Würthner et al., 2011). By contrast, MemBrigh-labeled EVs show an absorption spectrum identical to the solubilized form of the probe (Fig. 2B and supplementary Fig. 1A), revealing that the MemBright is efficiently embedded in EV membranes. The fluorescence spectra revealed that MemBright-labeled EVs are as bright as PKH-labeled EVs even though the MemBright was 10-fold less concentrated than PKH (Fig. 2B). This difference was confirmed by fluorescence microscopy (Fig. 2C) where we found that Zmel1 EVs labeled with MemBright (at 0,2μM) are as bright as identical EVs labeled with PKH (at 2μM) (Supplementary Fig. 2A). When both dyes were used at similar dilutions (0,2μM), the MemBright labeled EVs were much more bright than the PKH ones (Supplementary Fig. 2B). Indeed, MemBright displays >20-fold higher quantum yield than the PKH: 0.42 vs 0.02 (Table 7). Taking into account that MemBright-Cy3 and PKH26 contain the same Cy3-based fluorophore, such remarkable difference in the quantum yield suggests inefficient partitioning of PKH into EV membranes. This poor partitioning probably arises from aggregation of PKH in aqueous media, in line with characteristic short-wavelength shoulder in the absorption spectrum in the samples of EVs (Supplementary Fig. 1B). This is not the case for MemBright. Interestingly, a similar spectroscopic experiment conducted without EVs reveals the presence of a red-shifted fluorescence peak with PKH alone, but not with MemBright alone (Supplementary Fig. 1C). These fluorescent PKH aggregates have an average diameter of 80nm (± 10nm), as analyzed by Fluorescence Correlation Spectroscopy (FCS), which is in the range of EVs and therefore could lead to artifacts. These experiments reveal two clear advantages of MemBright *vs* PKH: (i) much more efficient staining of EVs with brighter fluorescence signal and (ii) non-fluorescence in the absence of EVs, unlike PKH showing emissive particles poorly distinguishable from EVs. To complement these studies and determine whether the MemBright could affect the morphology of labeled EVs, we analyzed them by electron microscopy. We found that neither the morphology, nor the size of MemBright-labeled EVs was affected, when compared to non-labelled EVs (Fig. 2D). Altogether, these experiments prove that labeling EVs with MemBright does not lead to soluble fluorescent aggregates that can be confounded with labeled EVs. In addition, given its high quantum yield, MemBright can be used at a relatively low concentration to efficiently label isolated EVs.

**Figure 2:**
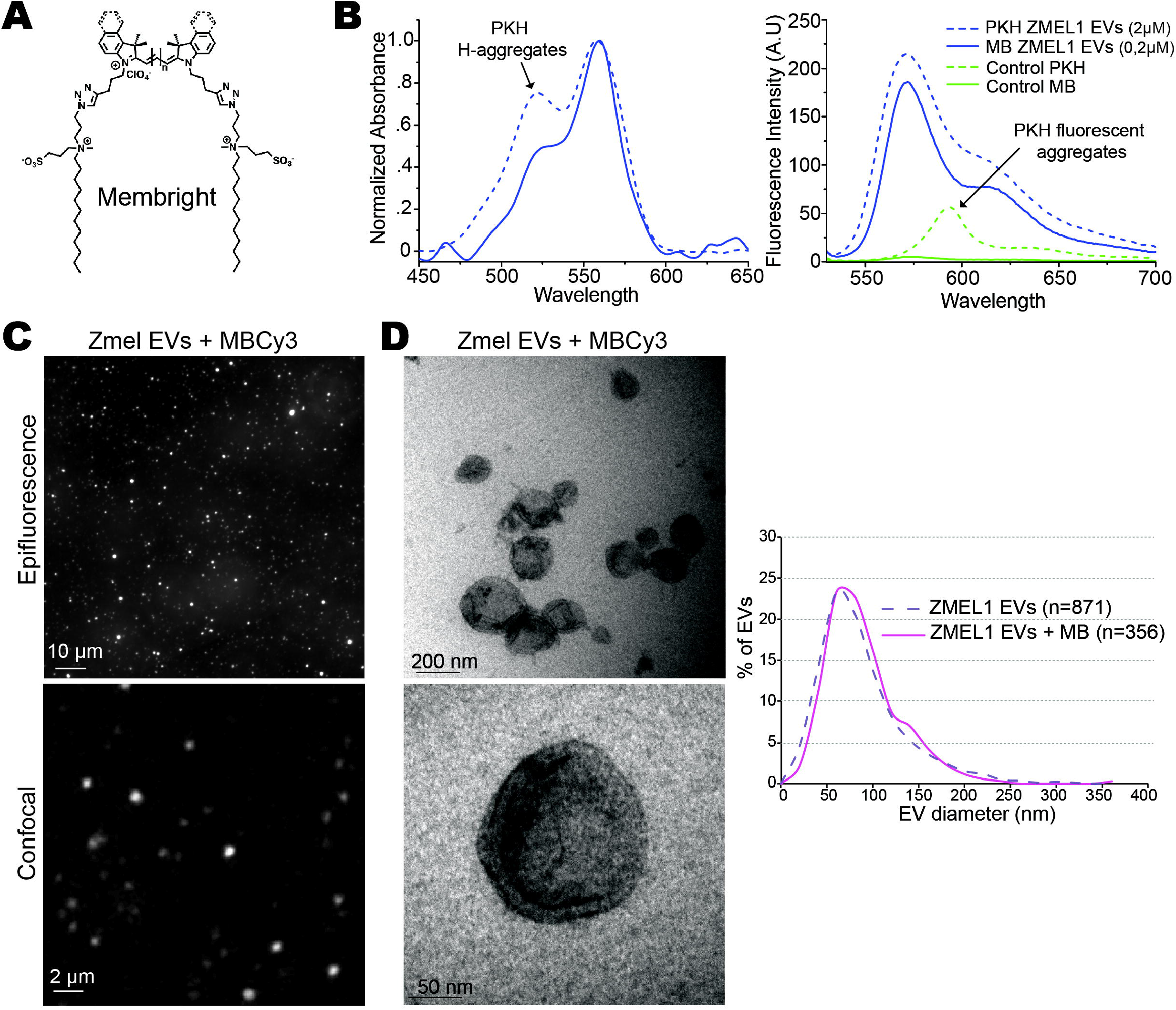
EVs can be brightly and specifically labeled with MemBright. **(A)** Molecular structure of the membrane binding probe MemBright. **(B)** Histograms showing a spectroscopy analysis of MemBright (MB) and PKH labeled Zmel1 EVs describing the absorbance (left histogram, y axis) and the fluorescence intensity (right histogram, y axis) versus the wavelength (nm, x axis). Arrows indicate the presence of PKH aggregates in labeled EVs (left) as well as in control PKH alone (right). **(C-D)** Representative images of Zmel EVs labeled with MemBright Cy3 (MBCy3) observed by Epifluorescence (upper) and confocal (lower) **(C)** or electron microscopy at low (upper) or high (lower) magnification and histogram showing the percentage of labeled and non-labeled Zmel1 EVs (y axis) versus their diameter (x axis, nm) by electron microscopy (right graph) **(D)**.

In order to know whether the MemBright can be used with EVs of different origins and perform complementary studies, we used EVs isolated from 4T1 mouse mammary carcinoma cells, where we could easily assess the presence of EVs using specific antibodies. Spectroscopy studies reveal that 4T1 EVs labeled with MemBright-Cy3 are brighter than 4T1 EVs labeled with PKH-26 and do not contain soluble fluorescent aggregates, unlike PKH-26 samples (Supplementary Fig. 1B and Table 7). Therefore, MemBright labels 4T1 EVs as well as Zmel1 EVs. In addition, electron microscopy showed that MemBright labeling does not affect the morphology or the diameter of 4T1 EVs (data not shown). Finally, we analyzed MemBright-labeled 4T1 EVs by density gradient and observed that the majority of the fluorescent MemBright is present in the fractions where most EVs are found, as confirmed by the presence of Alix and TSG101 (Supplementary Fig. 3). Altogether, our experiments demonstrate that MemBright does not lead to staining artifacts and offers a bright and specific labeling of EVs that is highly attractive for tracking EVs *in vivo*.

### Tumor EVs can be individually tracked in the living zebrafish embryo

We next investigated whether MemBright labeling could be used for tracking tumor EVs *in vivo*. We injected zebrafish embryos at 2 days post-fertilization with MemBright-labeled EVs in the duct of Cuvier, which connects the yolk to the heart and hence to the blood circulation. Using confocal microscopy, we observed the fluorescent EVs essentially in the tail region of the embryo, and not in the rest of the fish embryo. This region is composed of a main artery (named the dorsal aorta) and a complex venous network (named the caudal plexus) (Fig. 3A). Minutes following injection, we were able to observe several fluorescent EVs that were either still circulating in the blood flow or that were already arrested along the endothelium (Fig. 3A and Movie 1). Because we observed fluorescent objects of various sizes, we first assessed the apparent size of EVs by comparing them to 100nm fluorescent polystyrene beads. *In vitro*, we found that MemBright labeled EVs and 100nm fluorescent beads immobilized on surfaces display similar apparent sizes, which correspond to the resolution limits of confocal microscopy (Supplementary Fig.4A). Upon injection in the circulation of zebrafish embryos, we observed similar diffraction-limited spots for MemBright-labeled tumor EVs and the beads (Supplementary Fig.4B). These observations suggest that MemBright in combination with our microscopy set-up allow imaging of fluorescent objects of the size of an individual EV. At this stage, we can however not assess whether bigger spots result from bigger EVs or cluster of small EVs. In addition, MemBright can be used to co-inject different types of EVs labeled with different colors (Cy3, Cy5) and specifically track their fate. As a proof of concept, we co-injected Zmel1 tumor EVs (labeled with MemBright-Cy5) with 4T1 mouse tumor EVs (labeled with MemBright-Cy3) in embryos and observed both specific localizations for each EVs population as well as a common uptake in isolated cells (Supplementary Fig. 5). This suggests that MemBright could be used to follow specific internalization routes of distinct types of EVs that might be at the basis of their function and message delivery.

**Figure 3:**
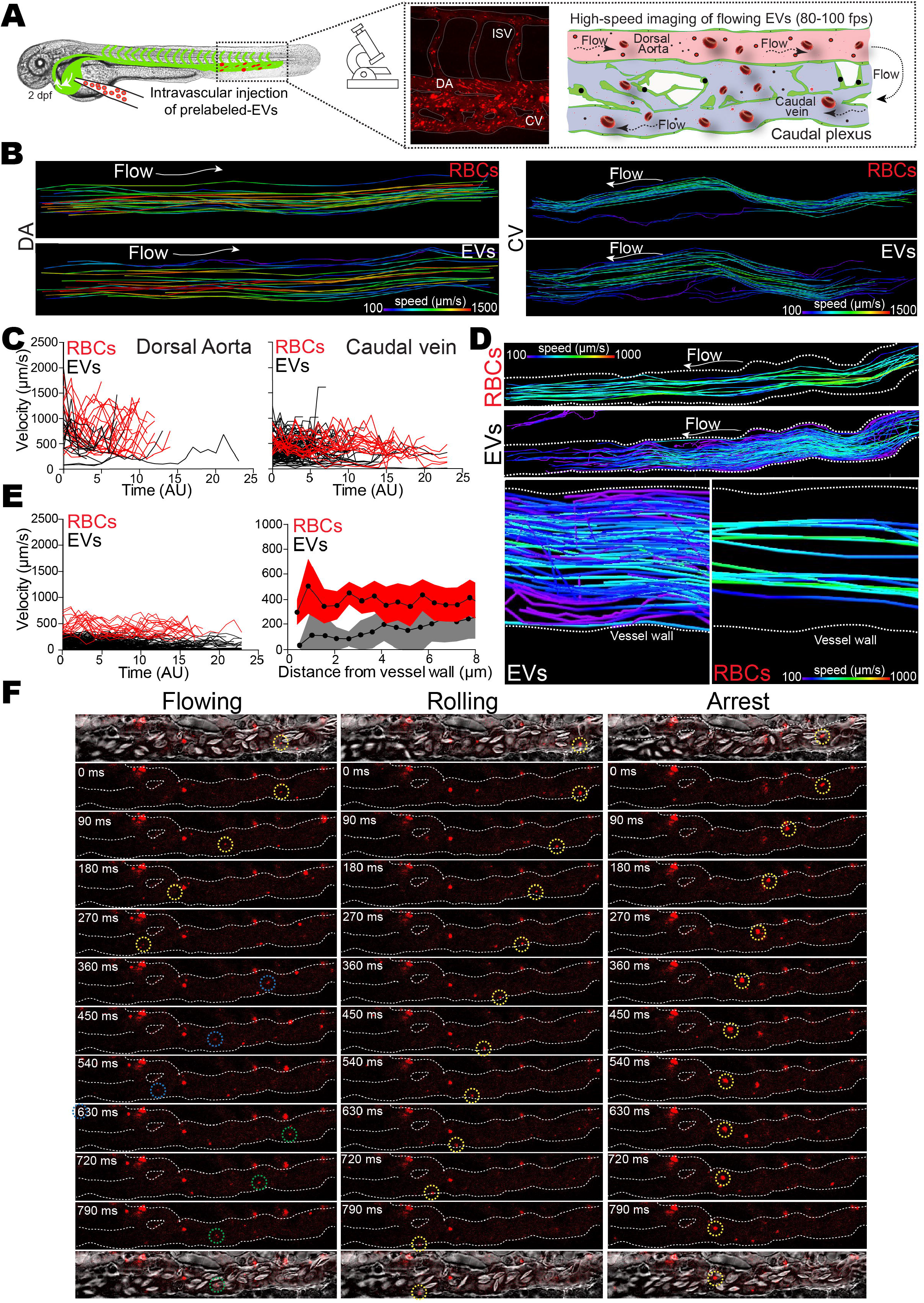
Hemodynamic characterization of individual EVs tracked in the circulation of zebrafish embryo. **(A)** Illustration of the experimental setup used to track circulating EVs: two days post-fertilization *Tg(Fli1:GFP)* zebrafish embryos were injected in the duct of Cuvier with MemBright-Cy3 labeled Zmel1 EVs (left) and observed in the caudal plexus with highspeed confocal microscopy (right). Middle: representative Z projection of MemBright-Cy3 Zmel1 EVs in the caudal plexus of a zebrafish embryo right after injection. Right: shematic representation of the caudal plexus showing the direction of the blood flow in the dorsal aorta (pink) and the veinous plexus (blue). **(B)** Individual tracks of red blood cells (RBC) or Zmel1 EVs in the dorsal aorta (DA, left) and in the caudal vein (CV, right). Color coding represents velocities. **(C)** Representative histograms showing the velocity (y axis, μm/s) versus the time (x axis, AU) of RBCs (red) and EVs (black) in the dorsal aorta (left) and in the caudal vein (right). **(D)** Upper: Individual tracks of red blood cells (RBC) or 4T1 EVs in the caudal vein. Lower: Zoom on individual tracks of red blood cells (RBC, right) or 4T1 EVs (left) in the caudal vein showing their proximity to the vessel wall (white lines). Color coding represents velocities. **(E)** Left: representative histogram showing the velocity (y axis, μm/s) versus the time (x axis, AU) of RBCs (red) and 4T1 EVs (black) in the caudal vein. Right: representative histogram showing the velocity (y axis, μm/s) versus the distance to the vessel wall (x axis, μm) of RBCs (red) and EVs (grey) in the caudal vein. **(F)** Examples of individual EVs flowing (left panels), rolling (middle panels) or arresting (right panels) in the circulation of the caudal vein.

We first aimed to describe the behavior of tumor EVs in the blood circulation, as it had never been characterized before. This challenging task requires a combined usage of a bright fluorescent signal with high sensitivity detection, which is a prerequisite for high-speed imaging. To do that, we performed high-speed confocal acquisitions of flowing tumor EVs (and of co-flowing red blood cells, RBCs) in different regions of the vasculature (dorsal aorta and caudal vein) of living zebrafish embryos (Fig. 3A and Movies 2 and 3). The motion of RBCs is used for probing flow profiles as well as potential differential hemodynamic behavior with tumor EVs, which significantly differ in size. When tracking both tumor EVs and RBCs and compiling hundreds of individual tracks, we first found that EVs have a higher velocity in the aorta than in the caudal veins, in accordance with the hydrodynamic profiles previously described in this region of the zebrafish embryo vasculature (Fig. 3B-C) (Follain et al., 2018a). Interestingly, tumor EVs tend to arrest in regions with lower flow velocities, such as the venous caudal plexus, suggesting that blood flow could control their probability of arrest (Fig. 4A). A similar observation has been done for endogenous EVs (Verweij *et al*., co-submitted). Second, when analyzing co-motion of tumor EVs and RBCs in a single vessel, we noticed that EVs have a reduced velocity compared to RBCs (Fig. 3C, E). To determine whether our observations were restricted to Zmel1 EVs, we injected MemBright-labeled 4T1 EVs (Fig. 3D). Similarly to Zmel1 EVs, 4T1 EVs display a higher velocity in the dorsal aorta than in the caudal veins, but a slower velocity than RBCs (Fig. 3E). Interestingly, we observed that the hemodynamic behavior of tumor EVs differs from the one of RBCs in regions close to the vessel wall, from which RBCs are mostly excluded. Indeed, when we plotted the velocity of tumor EVs as a function of their position in the vessel with regards to vessel walls, we observed that tumor EVs explore the vicinity of vessel walls with a reduced velocity (when compared to tumor EVs flowing in the center of the vessel) (Fig. 3D-E). It thus seems that tumor EVs follow a Poiseuille flow, which predicts that objects displaced by a laminar flow would have a reduced velocity, because of frictional forces, along the border of the vessel wall. Such a behavior, in addition to their potential adhesive capacity, could thus favor the arrest of tumor EVs. Indeed, individual inspection of EVs using our high-speed acquisitions reveals three distinct behaviors (Fig. 3F): i) Some tumor EVs flow in the center of the vessel or in the vicinity of the endothelium. In the center their velocity is similar to the velocity of RBCs, but when flowing close to the vessel wall, their velocity is decreased. ii) Some tumor EVs seem to roll on the surface of the endothelium. Their velocity is even lower than flowing tumor EVs and the rolling behavior suggests the involvement of adhesive molecules whose function would be favored by permissive flow profiles. iii) Some tumor EVs eventually arrest over the course of a few seconds of acquisition (Fig. 3F). We observed arrest of EVs following a rolling behavior, suggesting that it could be driven by rolling and activation of adhesion molecules, as well as sharp arrest of flowing EVs, without a rolling phase (Movie 4). A very similar behavior was observed for endogenous EVs (Verweij *et al*. co-submitted). Altogether, these observations suggest that the synergistic usage of a bright and specific dye, MemBright, with sensitive high-speed microscopy allows, for the first time, detection of flowing tumor EVs *in vivo* and dissection of their behavior in the vasculature.

**Figure 4:**
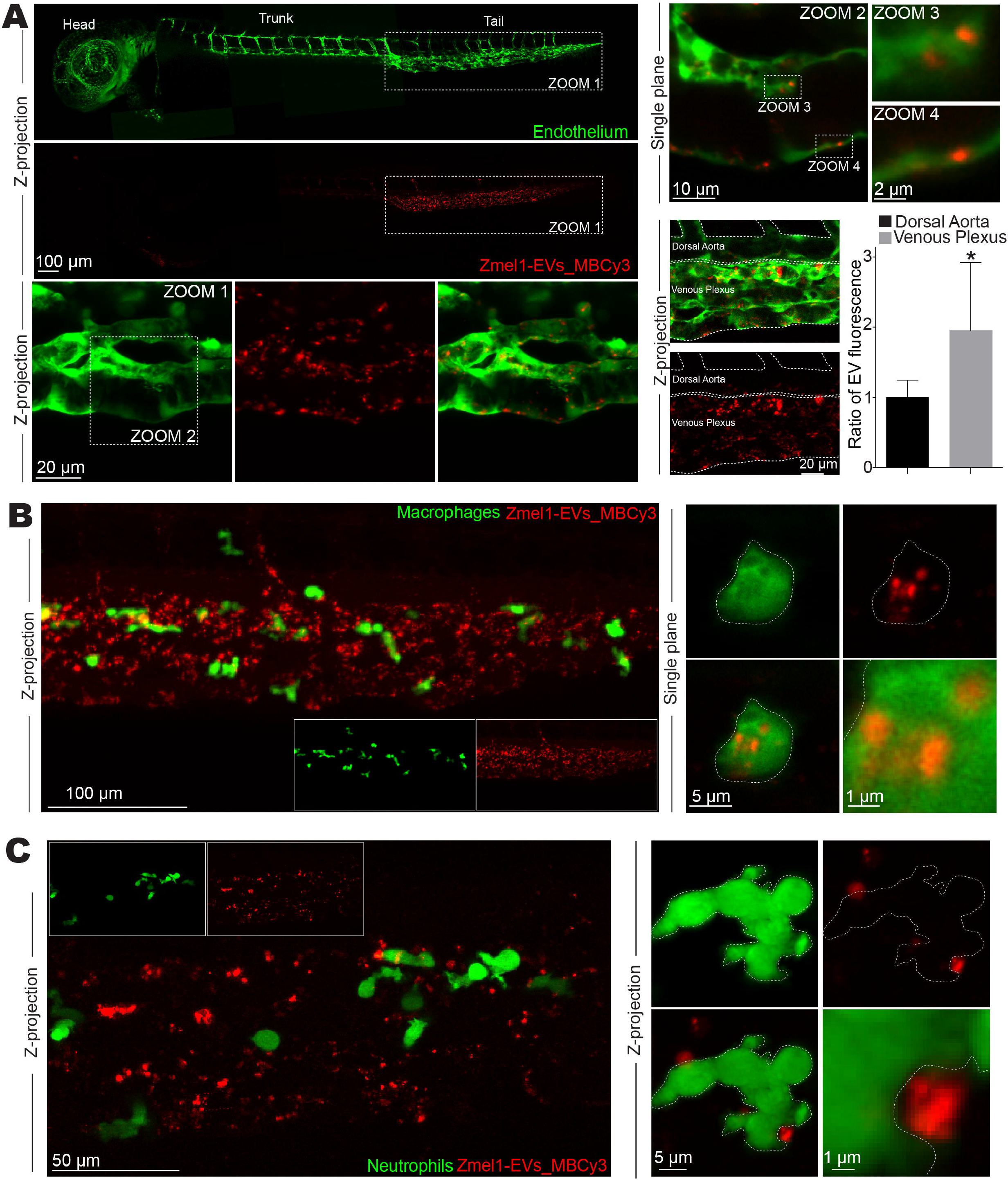
Zmel1 EVs are mainly uptaken by endothelial cells and macrophages, but not by neutrophils. **(A)** Representative confocal images of MemBright-Cy3 labeled Zmel1 EVs 3hours post injection in *Tg(Fli1:GFP)* embryos (endothelium specific expression) shown at different magnifications with Z projections (Left) or single plane (upper right). Lower right: Representative z-projections showing the borders of the dorsal aorta and the venous plexus and a histogram showing the EV fluorescence per surface in dorsal aorta and venous plexus (p<0,0001; Mann-Whitney test). **(B)** Representative confocal images of MemBright-Cy3 labeled Zmel1 EVs 3hours post injection in *Tg(mpeg1:GFP)* (macrophages specific expression) shown at low magnification with Z projections (Left) and high magnification with single plane (right). **(C)** Representative confocal images of MemBright-Cy3 labeled Zmel1 EVs 3hours post injection in *Tg(mpo1:GFP)* (neutrophils specific expression) shown at low magnification with Z projections (Left) and high magnification with single plane (right).

### Circulating tumor EVs are mostly uptaken by endothelial cells and patrolling macrophages

How circulating tumor EVs target specific cell types at distance remains a mystery, mostly because this step could not be captured before. Here, around 10 to 15 minutes following injection in the zebrafish embryo, most of the injected tumor EVs are found arrested, exclusively in the tail region of the fish (Fig. 4A). In addition, we found that most of the uptake by endothelial cells occurs in the venous region, rather than in the dorsal aorta (Fig. 4A) suggesting that the permissive flow profiles of this particular region favor arrest and uptake of tumor EVs, as they do for circulating tumor cells (Follain et al., 2018a). To assess which cell types could uptake tumor EVs, we used three transgenic zebrafish lines with different tissue specific GFP expression *(Tg(Fli1:GFP)* for the endothelium Fig. 4A, *Tg(mpeg1:GFP)* for macrophages Fig. 4B, and *Tg(mpo:GFP)* for neutrophils Fig. 4C). We found that tumor EVs are rapidly uptaken by endothelial and macrophages cells, but not by neutrophils (Fig. 4A, B and C) that are know to have a reduced phagocytic activity (Le Guyader et al., 2008). Embryos injected with the MemBright dye alone dot not show any signal that could arise from soluble fluorescent aggregates (Supplementary Fig. 6). We then generated a semi-automated method to precisely determine the percentage of EVs internalized by endothelial and macrophages 3 hours after injection. We found that both cell types uptake equivalent proportions of Zmel1 EVs, 43% (n= 18 fishes) and 38% (n=11) respectively. Together, this represents the large majority of arrested EVs in the zebrafish embryo at that stage. Importantly, Verweij and colleagues similarly show that endogenous CD63 positive EVs secreted by YSL accumulate specifically in endothelial cells and macrophages present in the caudal plexus and not in other cell types (Verweij *et al*. co-submitted), suggesting again that circulating EVs of different origins share common mechanisms of arrest *in vivo*. Different types of monocytes and macrophages have been previously shown to be able to uptake tumor EVs in mice (Whiteside, 2016). We noticed that tumor EVs are mostly uptaken by small and round mpeg1-positive cells present in the caudal plexus and not by more elongated ones (Fig. 5A-B). In non-injected embryos, these round cells are in direct contact with the blood flow (Fig. 5A) that they scan using long protrusions (Fig. 5C and Movie 5). Time lapses revealed that these small round macrophages have a reduced velocity compared to the elongated ones (Fig. 5D, Supplementary Fig. 7 and Movie 6). Therefore, the morphology, the location and the dynamics of these cells are reminiscent of patrolling monocytes, which are known to play an important role in tumor progression and metastasis in mice and men (Auffray et al., 2007; Carlin et al., 2013; Hanna et al., 2015). To confirm this observation and gain insight into the ultrastructure of these cells, we used our established Correlative Light and Electron Microscopy (CLEM) procedure (Goetz et al., 2014, 2015; Karreman et al., 2016b) on *Tg(mpeg1:GFP)* embryos injected with tumor EVs labeled with MemBright-Cy3 (Fig. 5E and Movie 7). We first imaged two typical mpeg1:GFP positive cells that have uptaken circulating tumor EVs using confocal microscopy in the living zebrafish embryo before engaging in the CLEM procedure. Once embryos were fixed and resin embedded, micro-computed tomography was used to narrow down the region of interest (ROI) as previously described (Karreman et al., 2016a). Serial transmission electron microscopy was performed and retrieval of the cell-containing region of interest was obtained through identification of anatomical landmarks (Supplementary Fig.8). Fine cellular segmentation allowed to create 3D models that revealed that macrophages localize in a cavity of the lumen of the vessel, where they form tight contacts with the endothelium and extend wide protrusions in the lumen (Fig. 5E and Movie 7). Interestingly, the region of the endothelium which is in contact with the macrophages is enriched of endocytic stuctures, suggesting active exchange between those two cell types (Fig. 5F). The macrophages that have uptaken tumor EVs extend long and dynamic protrusions in the lumen of the vessel (Fig. 5C and G), as it had been shown for patrolling monocytes in mice (Carlin et al., 2013). These protrusions are 3-4 micrometer long and their diameter is around 100nm, which makes them capable of scanning large portions of the vessel lumen. Surprisingly, analysis of the serial sections reveals that their height can be > 3 μm, and that these protrusions are actually forming large flat sheets deployed in the lumen. Altogether, our data show that circulating tumor EVs are rapidly uptaken by patrolling macrophages in the zebrafish embryo, which suggests that it can be used to track the mechanisms of delivery of tumor EVs at high spatio-temporal resolution.

**Figure 5:**
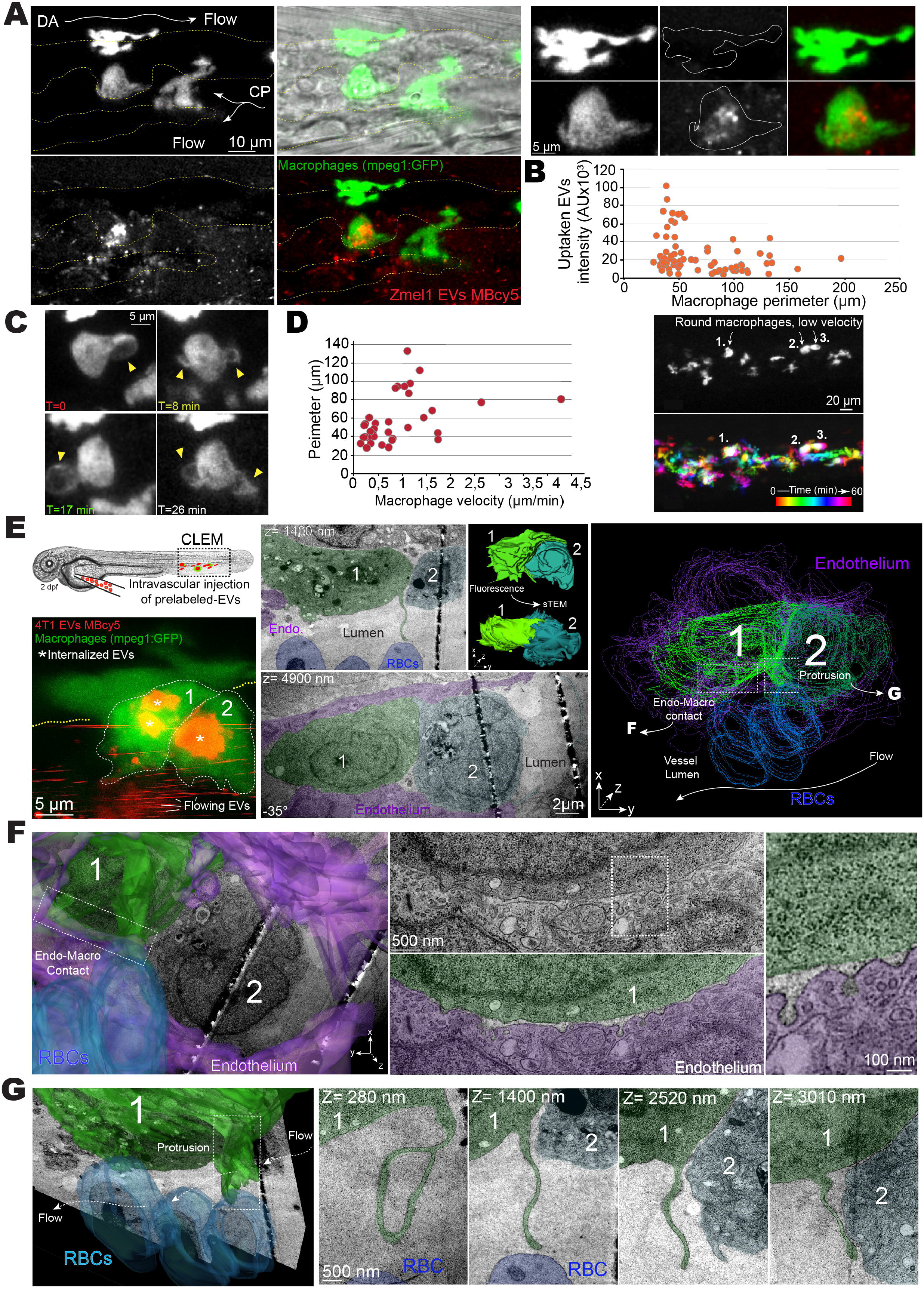
Circulating EVs are uptaken by patrolling macrophages. **(A)** Representative confocal Z projections images of MemBright-Cy3 labeled Zmel1 EVs injected in *Tg(mpeg1:GFP)* (macrophages specific expression) (left) with zooms on elongated macrophages devoid of EVs and round macrophages accumulating EVs (right). **(B)** Histogram showing the intensity of uptaken EVs (y axis, arbitrary units) versus the perimeter of the macrophages (x axis, μm). Each dot represents one macrophage. **(C)** Individual time points of single plane confocal images showing the dynamics of the protrusions in round macrophages. **(D)** Histogram showing the perimeter of macrophages (y axis, μm) versus their velocity (x axis, μm/s) (left) and images at the beginning (T=0) and the end (T=60min) of a representative time-lapse. Velocities of migration of *Tg(mpeg1:GFP)*-positive cells are represented with a color code. Three round *Tg(mpeg1:GFP)*-positive cells (1,2 and 3) show very little displacement during one hour. **(E)** CLEM experiment on *Tg(mpeg1:GFP)* embryos injected with MemBright-Cy3 4T1 EVs imaged by confocal right after injection (left, Z projection). Middle: electron microscopy images on two different Z planes showing the same cells. Right: 3D model showing the two macrophages (green), the endothelium (purple) and three red blood cells (blue). **(F)** Electron microscopy images of the contact between the endothelium and the macrophage, showing the accumulation of exocytic structures on the endothelium side. **(G)** 3D model and electron microscopy images of one protrusion sent by the macrophage into the lumen. This protrusion is visible over several microns in Z.

### Internalized tumor EVs are targeted to late endosomal compartments

To gain further insight into the mechanisms through which patrolling macrophages uptake tumor EVs, we assessed the dynamics of tumor EVs’ spreading in close proximity to target cells (Movies 8). We observed two different types of uptake. In the first one, EVs arrest at the surface of the macrophage and undergo a slow internalization that can be tracked at optimal spatio-temporal resolution (Fig. 6A and C; Movie 9). The timing of this uptake (~30sec) is in the range of classical endocytosis (Fig. 6A)(Idrissi and Geli, 2014; Taylor et al., 2011). We observed a second mechanism of internalization that is significantly faster (<5 sec). Here, tumor EVs are first caught by a protrusion extending from the macrophage, crawl back along this protrusion towards the cell centre before being internalized at the basis of the protrusion (Fig. 6B and C; Movie 10). At this stage, this mechanism is reminiscent of either filopodia surfing, a mechanism of EV uptake previously described *in vitro* (Heusermann et al., 2016) or of macropinocytosis, which has been shown to be responsible for the uptake of exosomes by microglia (Fitzner et al., 2011). Nevertheless, upon quantification of several similar uptakes, we confirmed that the latter mechanism is significantly faster than classical endocytosis (Fig.6C).

**Figure 6:**
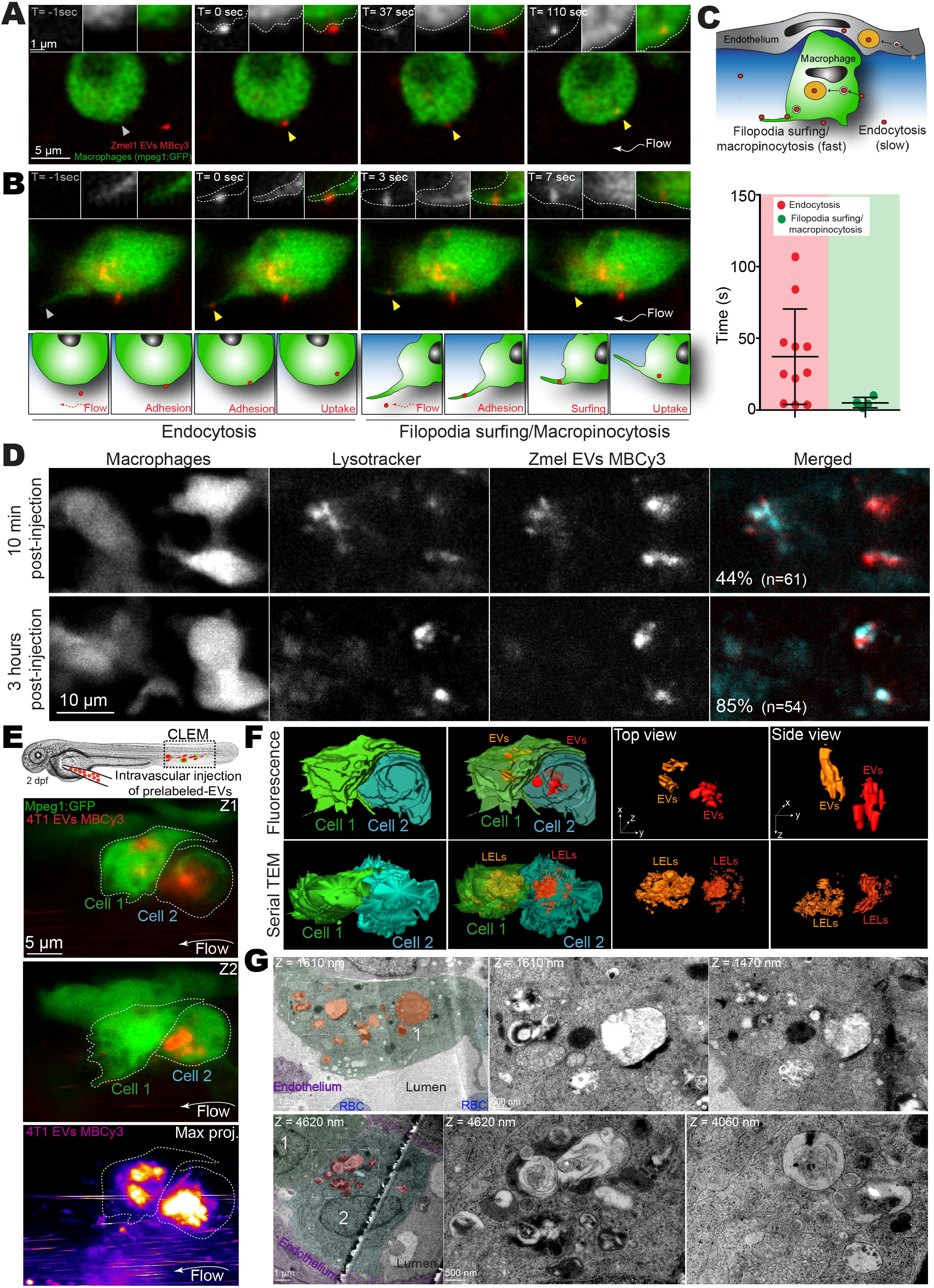
EVs are uptaken through different mechanisms and accumulate in late endosomal compartments. **(A)** Representative single plane confocal images of *Tg(mpeg1:GFP)* embryos injected with Zmel1 MemBright-Cy3 (MBCy3) EVs extracted from time-lapses generated right after injection and showing the attachment and uptake of EVs by endocytosis. **(B)** Representative single plane confocal images of *Tg(mpeg1:GFP)* embryos injected with Zmel1 MBCy3 EVs extracted from time-lapses generated right after injection and showing the sliding of EVs on the macrophage protrusion and its fast internalization. **(C)** Schematic representation of the modes of uptake by macrophages (upper) and histogram showing the duration (y axis, s) of those two mechanisms. **(D)** Representative confocal single planes of *Tg(mpeg1:GFP)* embryos injected with Zmel1 MBCy3 EVs and incubated with DeepRed lysotracker. **(E)** CLEM experiment on *Tg(mpeg1:GFP)* embryos injected with MemBright-Cy3 4T1 EVs imaged by confocal (single confocal planes of the GFP and the MBCy3 channels and Z projection of the EV chanel). **(F)** 3D model of the two cells and the uptaken EVs generated form the confocal data (upper panel, fluorescence) and 3D model of the two cells and the MVBs-late endosomes-lysosomes compartments (LELs) generated from the serial transmission electron microscopy data (lower panel, serial TEM). **(G)** Global view of each macrophage highlighting the MVBs-late endosomes-lysosomes compartments (orange and red, left). Zooms of those compartments are shown on the right in two different Z positions of the same region.

Next, we wondered which intracellular compartments are targeted by uptaken EVs. For this, we took advantage of the possibility to label late endosome-lysosomes by incubating the zebrafish embryo with the Lysotracker probe. *Tg*(*mpeg1:GFP*) embryos that were previously soaked in lysotracker were injected with MemBright-labeled Zmel1 EVs. We found that rapidly after injection, some MemBright labeled EVs are found co-localizing with lysotracker, although the majority is not (Fig.6D). Colocalization between these compartments and internalized tumor EVs increases over time and 3 hours post-injection, most tEV signal is found in endosome-lysosome compartments (Fig.6D). Of note, 24 hours post-injection, the MemBright signal is still visible and fully colocalizes with lysotracker. Although this approach provides a dynamic view of EVs trafficking in zebrafish embryos, lysotracker labeling does not distinguish between MVBs, late endosomes and lysosomes. To complement this study, we again exploited our established CLEM procedure on *Tg(mpeg1:GFP)* embryos injected with tumor EVs (Fig.5 E-G and Fig. 6E). We generated a 3D model of MemBright labeled EVs in each macrophage based on the confocal fluorescent data (called fluorescent 3D model, Fig. 6F, upper panel). In parallel, based on TEM serial sections of the same cells, we segmented all the MVBs, late endosomes and lysosomes that we could locate and generated a 3D model of these compartments (called TEM 3D model, LELs for Late Endosomes/Lysosomes) (Fig. 6F, lower panel and Movie 7). When comparing the two models, we found that the 3D model created from the fluorescent tumor EVs overlaps with the model from serial TEM sections of LELs (Movie 7). This suggests that the internalized tumor EVs are stored within these MVBs, late endosomes lysosomes compartments, that we imaged at high-resolution (Fig.6E, lower panels). Besides, close examination of the EM stack revealed EVs present in the lumen of the vessel, in close proximity of macrophage protrusions, as well as putative EVs present in endosomes (Supplementary Fig. 9). Altogether, this demonstrates the power of the zebrafish embryo to track, at multiple scales, the fate of nanometer-sized objects such as tumor EVs.

### Cells stained with MemBright can naturally release labeled exosomes *in vivo* and *in vitro*

We focused so far on tumor EVs that were previously isolated, labeled and tracked *in vivo*. This strategy, however, does not allow to track tumor EVs shed from *in vivo*-grown tumors. Interestingly, during the course of our experiments, we noticed that EVs can be labeled by incubating the secreting cells with the MemBright dye. We first observed that the MemBright exclusively accumulates in late endosomal compartments (identified by Lysotracker) only few minutes after its addition to Zmel1 cells in culture (Fig. 7A). We thus wondered whether such pre-labeling with MemBright would allow to track the release of fluorescent EVs in the culture medium. Therefore, we incubated cells with the MemBright dye for 20 minutes. Upon extensive washing and trypsinization to remove all traces of the dye, cells were seeded in new plates and finally grown for 24 hours (Fig. 7B). This experiment revealed the presence of fluorescently labeled EVs among EVs isolated from these cells (Fig. 7C). These EVs display a morphology and diameters similar to EVs from non-labeled cells, as revealed by electron microscopy (Fig. 7D). Since 20 minutes after its addition, the MemBright exclusively labels late endosomes and disappears from the plasma membrane, we postulated that these labeled EVs are exosomes and not microvesicles. To address this postulate, we labeled 4T1 cells expressing CD63-GFP with the MemBright (Supplementary Fig. 10A,B). As expected, MemBright completely colocalizes with CD63-GFP after 24 hours, showing that it fully localizes in late endosomes/lysosomes. We isolated EVs secreted by those cells and detected the presence of EVs positive for both MemBright and CD63-GFP in the 100 000g pellet, proving that the MemBright can label exosomes (Supplementary Fig. 10C). We observed puncta positive for CD63-GFP but not for MemBright and vice-versa. This suggests that the MemBright dye does not label all exosomes equally and illustrates the heterogeneity of exosomes, which has recently been described (Kowal et al., 2016). Altogether, these experiments suggest that the MemBright is rapidly endocytosed, targeted to MVBs and incorporated into the membrane of intra-luminal vesicles before being subsequently released outside of the cells attached to the membrane of exosomes. Such a behavior is extremely useful since it allows to label naturally released fluorescent EVs by pre-incubating cells with MemBright. To determine whether it could be used to track the transfer of EVs between cells, we first designed *in vitro* co-culture experiments. For this, we co-cultured Zmel1 pre-labeled with MemBright-Cy5 with Zmel1 cells expressing cytoplasmic tdTomato. After a week of co-culture, we observed several Cy5 fluorescent puncta accumulating in the cytoplasm of Zmel1 tdTomato cells, suggesting that indirectly labeled EVs successfully transfer between neighboring cells (Fig. 7E). Such a result opens the door to *in vivo* experiments where pre-labeled tumor cells would be grafted in zebrafish embryos (Fig. 7F, G).

**Figure 7:**
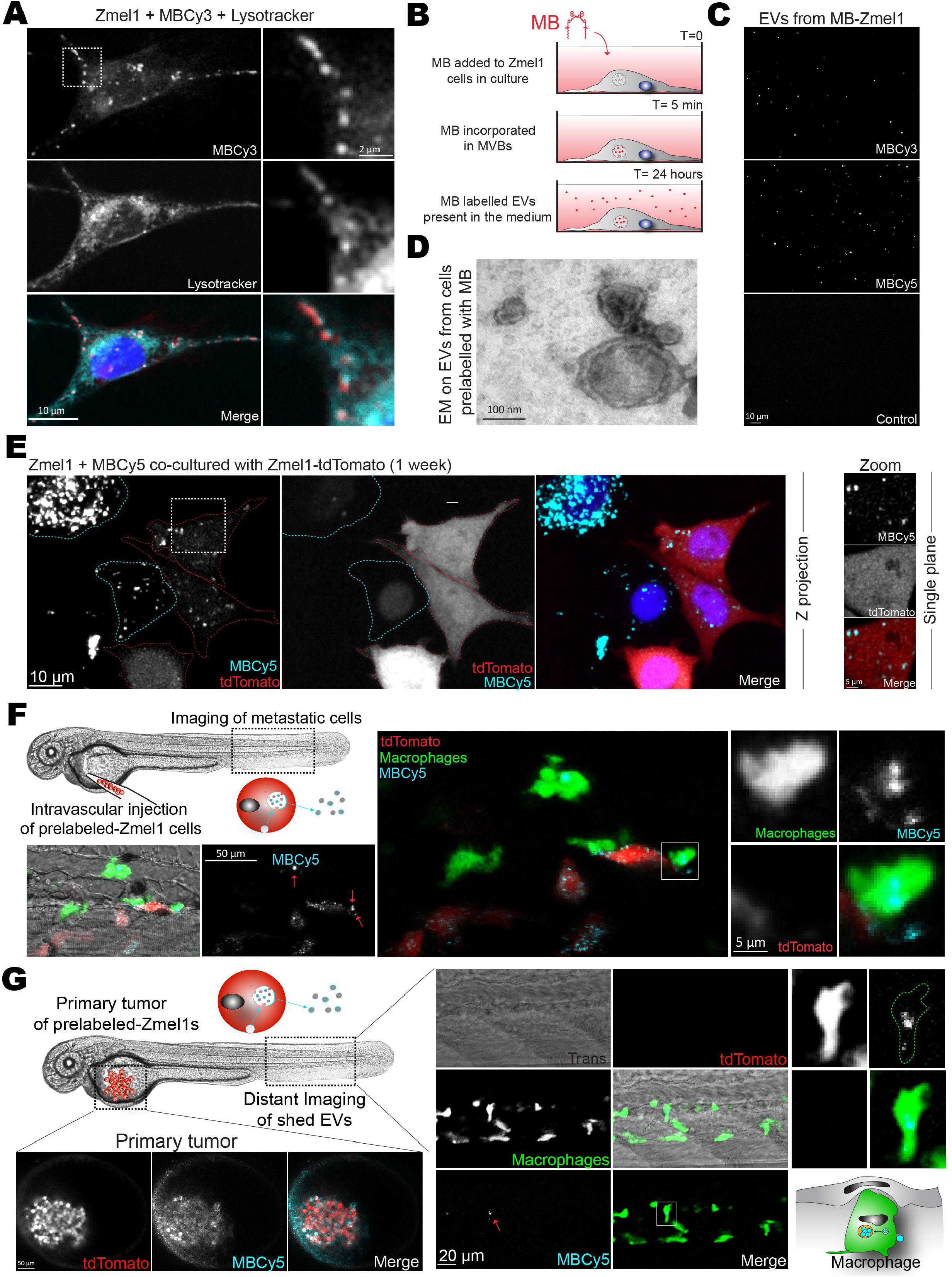
MemBright can be used to pre-label exosomes in secreting cells. **(A)** Representative confocal images of Zmel1 cells incubated with MemBright-Cy3 and stained with Lysotracker Deep Red. Right panel: Zoom showing colocalization between MemBright-Cy3 and Lysotracker Deep Red. **(B)** Schematic representation of the experimental procedure: MemBright added to cells in culture accumulates in MVBs and is subsequently released in exosomes **(C)** Representative images of EVs collected from Zmel1 cells labeled with MemBright-Cy3, MemBright-Cy5 or not labeled. **(D)** Electron microscopy image of EVs isolated from Zmel1 cells pre-labeled with MemBright-Cy3 showing their normal morphology. **(E)** Representative confocal images of Zmel1 cells prelabelled with MemBright-Cy5 and co-cultured with Zmel1 tdTomato cells, showing the transfer of MemBright in Z-projections (left) and single planes (right). **(F)** Confocal images of tdTomato Zmel1 cells prelabelled with MemBright-Cy5 injected in the circulation of *Tg(mpeg1:GFP)* embryos and imaged in the caudal plexus two days post-injection, showing the transfer of MemBright-Cy5 to macrophages. **(G)** Confocal images of tdTomato Zmel1 cells prelabelled with MemBright Cy5 injected above the yolk of *Tg(mpeg1:GFP)* embryos and imaged in the yolk region (primary tumor, left) and in the caudal plexus (distant imaging of shed EVs, right) two days post-injection, showing the transfer of MemBright-Cy5 to macrophages.

To probe local and distant transfer of EVs, we designed two types of experiments. To test local EVs transfer, tdTomato Zmel1 cells were pre-labeled with MemBright-Cy5 and subsequently injected into the circulation of *Tg(mpeg1:GFP)* zebrafish embryos. We have recently shown that tumor cells injected in the circulation of zebrafish embryos preferentially arrest and extravasate in specific regions of the caudal plexus (Follain et al., 2018a). When we imaged similar regions of embryos injected with pre-labeled Zmel1 cells, we observed many elongated myeloid cells, presumably macrophages crawling around arrested Zmel1 tumor cells (Movie 11). Importantly, we observed Cy5-positive fluorescent puncta present within these macrophages (Fig. 7F) suggesting local EVs transfer between tumor cells and macrophages. These puncta were not positive for tdTomato, revealing a different mechanism than the transfer of cytoplasmic material between melanoma cells and macrophages that has recently been documented in the zebrafish embryo (Roh-Johnson et al., 2017). In addition, we tested the distant transfer of EVs by injecting tdTomato Zmel1 cells that were pre-labeled with MemBrightcy5 in the yolk region and imaging macrophages present in the caudal plexus. Similarly to the previous experiment, we were able to detect Cy5 fluorescence in macrophages, suggesting the existence of a distant transfer of EVs that exploits the blood circulation for shedding and targeting at distance (Fig. 7G). Altogether, these experiments demonstrate that it is possible to use the MemBright to pre-label tumor cell exosomes and that this approach can be used both *in vivo* and *in vitro* to track the transfer of exosomes between tumor and stromal cells.

## Discussion

The work presented here establishes the zebrafish embryo as a new animal model to study tumor EVs *in vivo*. It demonstrates the proximity of zebrafish melanoma EVs to human melanoma EVs and shows how a novel membrane probe, the MemBright, specifically and brightly label EVs. Using this probe, we were able to precisely track the fate and behavior of EVs at high spatio-temporal resolution *in vivo*. This allowed us to provide the first description of the behavior of tumor EVs circulating in the blood flow, and to track their fate upon arrest. We identify the two main cell types uptaking circulating tumor EVs (endothelial cells and patrolling macrophages) and unravel their uptake mechanisms. Finally, we describe a novel method that exploits the properties of MemBright for pre-labeling of secreting cells to track transfer of exosomes *in vivo*. Such procedure could be widely used as an alternative method to cumbersome genetic encoding of exosome or EVs markers.

In a parallel study, Verweij and colleagues examine the fate of CD63 positive EVs secreted by the yolk syncytial layer in zebrafish embryo (Verweij *et al*. co-submitted). They track endogenous EVs, genetically labeled and naturally secreted during zebrafish development, while we tracked exogenous MemBright-labeled injected tumor EVs. Yet, both studies reach similar conclusions. They both show that: 1) endogenous and tumor EVs mainly arrest in the caudal plexus, in regions of low blood flow, 2) EVs are mostly uptaken by endothelial cells and patrolling macrophages, 3) EVs are stored in acidic compartments. Together, our reports establish the zebrafish embryo as a new model to study fundamental aspects of EVs biology *in vivo*. It thus represents a precious and complementary tool to invertebrate models *Drosophila* and *C. elegans*, which already contributed to better understand the mechanisms of EV secretion as well as their function (Beer and Wehman, 2017).

In addition, we propose the zebrafish embryo as a new and complementary model to murine and human cell culture systems for studying the fate and the function of tumor EVs during the priming of metastatic niches at distance. Compared to *in vitro* systems, zebrafish embryo offers an invaluable complex microenvironment, where different cell types known to contribute to tumor progression are present and can be tracked using established fluorescent transgenic lines. Its transparency allows to visualize individual tumor EVs dispersion and uptake in living zebrafish with unprecedented spatiotemporal resolution, which represents a major advantage over the mouse, where more complex intravital imaging procedures are required in order to visualize single EVs (Lai et al., 2015; Van Der Vos et al., 2016; Zomer et al., 2015). In addition to fast photonic imaging, the zebrafish embryo is amenable to CLEM, through procedures which are simplified compared to the mouse (Goetz et al., 2015; Karreman et al., 2016b). The possibility to switch from photonic to nanoscale electron microscopy using CLEM is perfectly suited to the study of small objects such as EVs. Such approach should allow to understand how tumor EVs secreted by a primary tumor reach the blood circulation before crossing the endothelium when reaching a given organ, but also to grasp the details of their uptake and trafficking at a subcellular level. For example, several studies performed with *in vitro* and mouse approaches reported the capacity of tumor EVs to cross and disrupt the endothelial barrier and hence promote metastasis (Tominaga et al., 2015; Treps et al., 2016; Zhou et al., 2014). Such behavior could thus be tracked at very high-resolution in the zebrafish embryo and, for example, highlight the disruption of the endothelial junctions by electron microscopy. Alternatively, CLEM could be used for tracking secretion mechanisms using methods described in Verweij *et al*. (Verweij et al. co-submitted) and thereby highlight how EVs are shed in the circulation, which is of utmost importance for their function.

Importantly, the complex yet stereotyped vascular network of the zebrafish embryo is easily accessible via conventional high-speed microscopy and thus offers fine description of circulating EVs behavior in the blood flow, which is not possible in any other animal model so far. This unique characteristic is crucial as 1) large amounts of circulating EVs are present in patient’s blood (Baran et al., 2010; Galindo-Hernandez et al., 2013; Logozzi et al., 2009) and 2) tumor EVs can travel through the blood circulation to modify the microenvironment in organs distant from the primary tumor (Costa-Silva et al., 2015; Grange et al., 2011; Hoshino et al., 2015; Liu et al., 2016; Peinado et al., 2012). Our study demonstrates that functional tests aiming to address the contribution of injected tumor EVs to tumor growth, extravasation and metastasis in zebrafish are at reach. Interestingly, the mechanisms by which circulating tumor EVs arrest in specific vascular regions and target residing cells are only poorly understood. Thanks to their relative low size, tumor EVs bear potentially margination properties that favor their intra-circulation transit along the vessel walls (Müller et al., 2014a), while red blood cells travel in the center of the vessel because of the hydrodynamic interactions (or lift force) with the walls (Cantat and Misbah, 1999). When analyzing the radial distribution of circulating tumor EVs in the circulation of zebrafish embryos, we observed that tumor EVs displayed a uniform density over the cross section, with velocities that were significantly lower in near-wall regions. While such property is currently investigated to foster delivery of drug-loaded nano- and microparticles (Lee et al., 2013), it demonstrates that the hemodynamic behavior of tumor EVs, that is inherent to their low size, might potentiate their ability to reach specific vascular regions. Interestingly, small particles (<100 nm in diameter) have been shown to exhibit minimal particle margination (Lee et al., 2013) that can explain long-lasting systemic circulation in the center of blood vessel with the RBC core, with no attachment to the vessel wall (van der Meel et al., 2014). Further high-spatiotemporal analysis of tumor EVs with controlled size could thus provide very helpful insight on their ability to target and arrest at specific vascular regions.

The potency of the zebrafish embryo and its associated methods is illustrated by our fine description of the uptake of a large proportion of circulating tumor EVs by a subset of macrophages. We consider these cells as functionally similar to murine and human patrolling monocytes for the following reasons: *1)* they are positive for the mpeg1 promoter (Ellett et al., 2011) which is expressed both by monocytes and macrophages in human (Karlsson et al., 2008; Spilsbury et al., 1995) *2)* they are small, round and have a slow migration velocity when compared to elongated differentiated macrophages *3)* they are sending highly dynamic protrusions toward the lumen of the vessels and show areas of direct cell-cell contacts with the endothelial wall, as previously shown (Colucci-Guyon et al., 2011; Murayama et al., 2006). These two last aspects are matching the main characteristics of human and mice patrolling monocytes (Auffray et al., 2007; Carlin et al., 2013). CLEM analysis of two patrolling macrophages that internalized circulating tumor EVs highlights two of these aspects. First, we describe large regions of contact between endothelial cells and macrophages. Such regions are enriched in endocytic structures, essentially on the endothelial side, and suggest active exchange that occurs at the border of these two cell types. Second, volume EM analysis reveals that the dynamic protrusions observed in live imaging are actually flat sheets of several microns that scan the vessel lumen. These structures could function as butterfly nets to catch small objects circulating in the lumen such as tumor EVs. Their length is very likely fostering the efficiency of trapping circulating objects very deep in the vessel lumen. Previous work in zebrafish had shown that such structures are specific to macrophages, alllowing them to internalize fluid-borne objects, unlike neutrophils that only phagocytose surface-bound ones (Colucci-Guyon et al., 2011). Once they have contacted the protrusion, the EVs quickly slide toward the cell body through unknown mechanisms, which could be similar to the filopodia surfing recently described (Heusermann et al., 2016). Those protrusions could also participate in macropinocytic uptake of EVs, similar to what has been observed by microglia (Fitzner et al., 2011). EVs are then internalized at the basis of the protrusions, probably in regions of active endocytosis (see Supplementary Fig. 9). Interestingly, our EM data revealed several EVs present at the basis of protrusions (see supplemental figure 9). Alternatively to this protrusion based catching mechanism, circulating EVs can directly bind to the macrophage. They can remain associated with the surface of the macrophage for a longer time, before being endocytosed. The capacity of patrolling monocytes/macrophages to rapidly uptake circulating EVs explains the very short half-life (10-20 minutes) of circulating EVs after their injection in either mouse (Morishita et al., 2015; Saunderson et al., 2014; Takahashi et al., 2013) or fish (our work) blood circulation. This is in agreement with the observation that chemical depletion of monocytes and macrophages in mice dramatically increases the stability of circulating EVs (Imai et al., 2015). In addition, we observed positive transfer between pre-labeled melanomas cells that have just extravasated and macrophages. This suggests that a direct transfer of tumor EVs to macrophages is likely to occur between cells in close proximity. Such transfer could recruit macrophages to tumor cells, which have recently been suggested to foster melanoma invasion upon direct transfer of material (Roh-Johnson et al., 2017).

Our fluorescence and electron microscopy analysis reveal that after their uptake, tumor EVs are stored in acidic degradative compartments, similarly to what has been described for macrophages *in vitro* (Feng et al., 2010). Determining whether and how uptaken EVs deliver a signal to the receiving cell, although they are mostly targeted to degradative compartments, is a central question in the EV field. It will be particularly important to address it in the case of tumor EVs uptaken by patrolling macrophages. Indeed, and by contrast to more differentiated tumor associated macrophages (Engblom et al., 2016), patrolling macrophages prevent tumor growth and metastasis (Hanna et al., 2015), and this capacity has recently been shown to involve tumor EVs (Plebanek et al., 2017). It is interesting to note that uptake mechanisms and compartments are similar between exogenous tumor EVs (this study) and endogenous EVs (Verweij *et al*. co-submitted). This suggests that tumor EVs are internalized using universal mechanisms and further demonstrates that the zebrafish embryo is a perfect model for dissecting such behavior.

Finally, the zebrafish embryo allows a direct comparison of EVs with distinct sizes, contents or origins. This will be essential to better understand the heterogeneity of EVs, as it is now clear that multiple sub-populations (or sizes) of EVs co-exist with different cargo contents and presumably different functions (Kowal et al., 2016). Co-injection of different types of EVs can, for instance, be used to precisely dissect the involvement of one given EV transmembrane or cargo protein, or to compare tumor EVs from patients at different stages of tumor progression. Using multi-color MemBright probes (Cy3, 5 or 7) to label EVs, it is possible to directly compare the behavior of co-injected populations of EVs. Labeling EVs with membrane probes after their isolation is fast and allows obtaining bright fluorescent EVs regardless of their origin. It is particularly relevant for EVs isolated from cell lines reluctant to gene expression manipulation, from animal body fluids, or, importantly in the case of tumor EVs from samples of cancer patients. However, the use of membrane probes requires the assurance of labeling specificity. This is particularly essential for studies aiming to track EVs dispersion and uptake, as dye aggregates can easily be confounded with EVs, due to their small sizes (Lai et al., 2015; Takov et al., 2017). Here, using spectroscopic and microscopic approaches, we have shown that the MemBright does not form such fluorescent aggregates, in contrast to commonly used PKH. In addition, it is brighter and can therefore be used at reduced concentrations, minimizing again the risk of false positive results. The key difference of MemBright from PKH is the presence of amphiphilic groups, which favor efficient transfer of the fluorophore from aqueous media to lipid membranes (Collot et al., 2015; Kucherak et al., 2010). Therefore, MemBright can be used to confidently track EV dispersion and uptake.

Altogether, our work on the tracking of exogenous tumor EVs (this study) and of endogenous EVs (Verweij *et al*. co-submitted) set the zebrafish embryo as a new and highly attractive *in vivo* model to track EVs at the single EV scale. Interestingly, both studies identified similar mechanisms of transit and uptake for physiological and pathological extracellular vesicles, which further validate the zebrafish embryo as a reliable animal model for studying the biology of EVs. Finally, we believe that the zebrafish embryo will open new avenues for EV biology as it offers adapted time and space scales to the study of small organelles *in vivo*.

## METHODS

### Cell culture and 4T1 CD63-GFP cell generation

Zmel1 cells and Zmel1 cells stably expressing cytoplasmic tdTomato, were kindly provided by Richard White (Memorial Sloan Kettering Cancer Center, New York). They were grown as previously described (Heilmann et al., 2015), in DMEM with 4.5 g/l glucose (Dutscher) supplemented with 10% FBS, 1% NEAA and 1% penicillin-streptomycin (Gibco) at 28°C, 5% CO_2_. All mammalian cells were grown at 37°C, 5% CO_2_. Mouse melanoma cell lines B16-F0, F1 and F10 were purchased from ATCC. They were grown in DMEM supplemented with 10% (v/v) EV-depleted fetal bovine serum (EV-d-FBS), glutamine 2mM and gentamicin. Mouse melanocyte cell line melan-a was kindly provided by Dr. Dorothy C. Bennett, (St. George’s University of London), and grown in RPMI, fetal calf serum 10%, TPA (200 nM). Primary melanocyte cultures and human melanoma cells were kindly given by Dr. M. Soengas (CNIO, Madrid). Melanocytes were cultured in 254CF medium (Gibco) supplemented with calcium chloride and HMGS (Gibco) and human melanoma cells were cultured in DMEM with 10% EV-d-FBS. Mouse mammary carcinoma 4T1 cells were grown in RPMI with 10% FBS and 1% penicillin-streptomycin (Gibco). 4T1 cells expressing CD63-GFP were generated as follows. Briefly, human CD63 cDNA was fused to AcGFP cDNA by In-Fusion cloning (Takara, Ozyme, Saint-Quentin-en-Yvelines, France) and introduced in pLenti CMV-MABBXXS mPGK-Blast vector. Lentiviruses were obtained by HEK293T cells (ATCC CRL-3216; cultured in DMEM, 10% FCS, 1% penicillin-streptomycin) transfection (Invitrogen, Life Technologies, Saint Aubin, France) with pLenti CMV-CD63-acGFP mPGK-Blast together with pLP1, pLP2 and pLP/VSVG lentiviral packaging plasmids to obtain lentiviral particles. After 48 hours, conditioned media was collected, filtered through a 0.22 μm filter to remove cell debris, and used to transduce 4T1 cells cultured in DMEM supplemented with 10% fetal calf serum and 1% penicillin-streptomycin (Gibco, USA) in the presence of 5μg/mL polybrene (Sigma Aldrich, Lyon, France), followed by selection with puromycin (1 μg/mL, Sigma Aldrich, Lyon, France).

### EV isolation and analysis

For Zmel1 and 4T1 EVs isolation, cells were cultured in EV depleted medium (obtained by overnight ultracentrifugation at 100,000g, using a Beckman, XL-70 centrifuge with a Ti70 rotor) for 24h before supernatant collection. Extracellular medium was concentrated using a Centricon Plus-70 centifugal filter (10k; Millipore) and EVs were isolated by successive centrifugation at 4°C: 5 minutes at 300 g, 10 minutes at 2,000 g, 30 minutes at 10,000 g and 70 minutes at 100,000 g (using a Beckman XL-70 centrifuge with a SW28 rotor). EVs pellets were washed in PBS, centrifuged again at 100,000 g for 70 minutes, resuspended in PBS and stored at 4°C. For *in vivo* experiments, EVs were used immediately after isolation or kept 4°C at and used the next day.

For mouse and human melanoma and melanocyte EVs isolation, cells were cultured in media supplemented with 10% EV-depleted FBS (FBS, Hyclone). FBS was depleted of bovine EVs by ultracentrifugation at 100,000xg for 70 min. EVs were isolated from conditioned media collected after 72 h of cell cultures by successive centrifugation at 10°C: 5 minutes at 300 g, 10 minutes at 500 g, 20 minutes at 12,000 g and 70 minutes at 100,000 g (using a Beckman Optima X100 with a Beckman 70Ti rotor). EVs pellets were washed in PBS, centrifuged again at 100,000 g for 70 minutes, and resuspended in PBS. Protein content was measured by bicinchoninic acid assay (BCA assay).

For transmitted electron microscopy analysis, 3 μl of EV extracts were allowed to dry on formvar coated grids for 20 minutes, fixed in 3% PFA for 10 minutes, rinsed in water and contrasted in a uranyl acetate (0,4%)/ methylcellulose (2%) mix for 10 minutes on ice. EVs were observed either with an Orius 100 charge-coupled device camera (Gatan) mounted on a Philips CM12 microscope operated at 80kV or with a Veleta 2kx2k side-mounted TEM CDD Camera (Olympus Soft Imaging Solutions) mounted on a Philips CM120 microscope operated at 120kV.

NTA was performed on Zmel1 EVs diluted 10 times with sterile PBS, using a Nanosight NS300 (Malvern Instruments). Measurement was repeated three times.

For density gradient analysis, EVs isolated in the 100,000 g pellet were loaded on top of a 5-40% iodixanol (optiprep) density gradient prepared as previously described (Deun et al., 2014). The gradient was centrifuged for 18 hours at 100,000g and 4°C (using a Beckman XL-70 centrifuge with a SW28 rotor). Gradient fractions of 1ml were collected from the top of the gradient. Fractions 1 to 4, 5 to 10 and 11 to 16 were pooled, diluted to 16 ml in PBS and centrifuged for 3 hours at 100,000g and 4°C. The resulting pellet was resuspended in 50 μl of PBS. For western blotting analysis, 10 μl of EV extracts were loaded on 4-20% polyacrylamide gels (Biorad), under denaturing conditions. The following antibodies were used: Alix (BD Biosciences 611621) and TSG101 (GeneTex GTX70255). Acquisitions were done using a PXi system (Syngene).

### Mass spectrometry analysis of Zmel1 EVs

#### Sample preparation

Zmel1 EV protein concentration was measured (RC-DC^™^; Bio-Rad, Hercules, CA) and 20-μg of EVs were denatured at 95°C for 5 min in Laemmli buffer, and then concentrated in one stacking band using a 5% SDS-PAGE gel. The gel was fixed with 50% ethanol/3% phosphoric acid and stained with colloidal Coomassie Brilliant Blue. Each band was excised, cut in five pieces, and transferred into a 96-wells microtiter plate. Gel slices were washed with 3 cycles of incubations in 100 μL of 50:50 (v/v) 25 mM NH_4_HCO_3_/ACN for 10 min. Gel bands were dehydrated with 50 μL 100% ACN and then reduced with 50 μL 10 mM DTT for 30 min at 60°C, followed by 30 min at RT. Proteins were then alkylated with 50 μL 55 mM iodoacetamide for 20 min in the dark at RT, and then 100 μL ACN was added for 5 min. Samples were washed with 50 μL 25 mM NH_4_HCO_3_ for 10 min, and then 50 μL ACN for 5 min, before being dehydrated with two cycles of incubations in 50 μL ACN for 5 min. Proteins were digested overnight with a modified porcine trypsin (Promega, Madison, WI) solution at a 1:100 (w/w) enzyme/protein ratio at 37°C. Tryptic peptides were extracted under agitation at RT with 60 μL 60% ACN/0.1% FA for 45 min, and then 100% ACN for 10 min. The extraction supernatants were pooled and vacuum-dried, before re-suspension in 40 μL 2% ACN/0.1% FA.

<u>Nano-LC-MS/MS analysis</u> was performed on a nanoAcquity UPLC device (Waters, Milford, MA) coupled to a Q-Exactive Plus mass spectrometer (Thermo Fisher Scientific, Bremen, Germany). The solvents consisted of 0.1% FA in H_2_O (solvent A) and 0.1% in ACN (solvent B). 1 μL of samples was loaded onto a Symmetry C18 precolumn (20 mm × 180 μm, 5 μm diameter particles; Waters, Milford, MA) over 3 min at 5 μL/min with 1% solvent B. Peptides were eluted on a Acquity UPLC BEH130 C18 column (250 mm × 75 μm, 1.7 μm particles; Waters, Milford, MA) at 450 μL/min with the following gradient of solvent B: linear from 1% to 8 % in 2 min, linear from 8% to 35% in 77 min, linear from 35% to 90% in 1 min, isocratic at 90% for 5 min, down to 1% in 2 min and isocratic at 1% for 2 min.

The Q-Exactive Plus was operated in data-dependent acquisition mode by automatically switching between full MS and consecutive MS/MS acquisitions. Full-scan MS spectra were collected from 300-1,800 m/z at a resolution of 70,000 at 200 m/z with an automatic gain control target fixed at 3 × 10^6^ ions and a maximum injection time of 50 ms. The top 10 precursor ions with an intensity exceeding 2 × 10^5^ ions and charge states ≥ 2 were selected on each MS spectrum for fragmentation by higher-energy collisional dissociation. MS/MS spectra were collected at a resolution of 17,500 at 200 m/z with a fixed first mass at 100 m/z, an automatic gain control target fixed at 1 × 10^5^ ions and a maximum injection time of 100 ms. A dynamic exclusion time was set to 60 s.

#### Data interpretation

MS/MS data were searched using Mascot^™^ (version 2.5.1, Matrix Science, London, UK) against an in-house concatenated target-decoy *Danio rerio-Bos taurus* UniProtKB database (91 038 entries, release 11/2016), containing trypsin and known contaminants, that was generated with the database toolbox from MSDA (Carapito et al., 2014). The following parameters were applied: a mass tolerance of 5 ppm on the precursor ions and 0.07 Da on the peptide fragments, oxidation of methionine residues and carbamidomethylation of cysteine residues as variable modifications, one permitted missed cleavage per peptide. Mascot result files were loaded into Proline software (http://proline.profiproteomics.fr/; (Carapito et al., 2015)) and proteins were validated on pretty rank equal to 1, 1% FDR on peptide spectrum matches based on adjusted e-value, at least 1 specific peptide per protein, and 1% FDR on protein sets based on Mascot Modified Mudpit scoring.

### Mass spectrometry analysis of mammalian EVs

Samples were lysed in urea and digested with Lys-C/trypsin using the standard FASP protocol. Peptides were analyzed by LC-MS/MS analysis using a LTQ Orbitrap Velos (Thermo Fisher Scientific). Raw files were analyzed with MaxQuant against a human protein database and the MaxLFQ algorithm was used for label-free protein quantification.

#### Sample preparation

Proteins were solubilized using 8 M urea in 100 mM Tris-HCl pH 8.0. Samples (7.5 μg) were digested by means of the standard FASP protocol. Briefly, proteins were reduced (10 mM DTT, 30 min, RT), alkylated (55 mM IA, 20 min in the dark, RT) and sequentially digested with Lys-C (Wako) (protein:enzyme ratio 1:50, o/n at RT) and trypsin (Promega) (protein:enzyme ratio 1:100, 6 h at 37 ° C). Resulting peptides were desalted using C_18_ stage-tips.

#### Nano-LC-MS/MS analysis

LC-MS/MS was done by coupling a nanoLC-Ultra 1D+ system (Eksigent) to a LTQ Orbitrap Velos mass spectrometer (Thermo Fisher Scientific) via a Nanospray Flex source (Thermo Fisher Scientific). Peptides were loaded into a trap column (NS-MP-10 BioSphere C18 5 μm, 20 mm length, Nanoseparations) for 10 min at a flow rate of 2.5 μl/min in 0.1% FA. Then peptides were transferred to an analytical column (ReproSil Pur C18-AQ 2.4 μm, 500 mm length and 0.075 mm ID) and separated using a 120 min linear gradient (buffer A: 4% ACN, 0.1% FA; buffer B: 100% ACN, 0.1% FA) at a flow rate of 250 nL/min. The gradient used was: 0-2 min 6% B, 2-103 min 30% B, 103-113 min 98% B, 113-120 min 2% B. The peptides were electrosprayed (1.8 kV) into the mass spectrometer with a PicoTip emitter (360/20 Tube OD/ID μm, tip ID 10 μm) (New Objective), a heated capillary temperature of 325°C and S-Lens RF level of 60%. The mass spectrometer was operated in a data-dependent mode, with an automatic switch between MS and MS/MS scans using a top 15 method (threshold signal ≥ 800 counts and dynamic exclusion of 60 sec). MS spectra (350-1500 m/z) were acquired in the Orbitrap with a resolution of 60,000 FWHM (400 m/z). Peptides were isolated using a 1.5 Th window and fragmented using collision induced dissociation (CID) with linear ion trap read out at a NCE of 35% (0.25 Q-value and 10 ms activation time). The ion target values were 1E6 for MS (500 ms max injection time) and 5000 for MS/MS (100 ms max injection time).

#### Data interpretation

Raw files were processed with MaxQuant (v 1.5.1.2) using the standard settings against a human protein database (UniProtKB/Swiss-Prot, August 2014, 20,187 sequences) supplemented with contaminants. Label-free quantification was done with match between runs (match window of 0.7 min and alignment window of 20 min). Carbamidomethylation of cysteines was set as a fixed modification whereas oxidation of methionines and protein N-term acetylation as variable modifications. Minimal peptide length was set to 7 amino acids and a maximum of two tryptic missed-cleavages were allowed.

### Protein comparisons

To compare the Zmel1 protein content with mammalian EV content, each protein list was concatenated and duplicate proteins were deleted. Ortholog proteins were searched using the ortholog protein files predicted by the PANTHER classification system (ftp://ftp.pantherdb.org/ortholog/13.0/ (Thomas et al., 2003)). Only proteins referred as “Least diverged ortholog” or “Ortholog” were considered. All comparisons between Zmel1 EVs and mammalian EVs were done using human orthologs and the lists of common proteins was obtained using Venny 2.1 (Oliveros, 2007).

### MemBright and PKH labeling of EVs

Isolated EVs were incubated with MemBright-Cy3 or Cy5 at 200nM (final concentration) in PBS for 30 minutes at room temperature in the dark. They were then rinsed in 15ml of PBS and centrifuged at 100,000g with a SW28 rotor in a Beckman XL-70 centrifuge. Pellets were resuspended in 50 μl PBS and stored at 4°C. For *in vivo* experiments, EVs were used immediately after isolation or stored overnight at 4°C and injected the next day. For PKH-26 labelling EVs were treated according to the manufacturer’s instructions (2 μM final concentration). Briefly, EVs in 200 μl of PBS were first mixed with 300 μl of Diluent C, then with 500μl of Diluent C containing 4 μl of PKH and finally incubated for 30 minutes at room temperature in the dark. PKH labelled EVs were then processed as MemBright labelled EVs. As control, PBS alone was processed similarly to EVs, labelled with MemBright or PKH and analysed by microscopy or spectroscopy.

For photonic microscopy analysis, 3 μl of labelled EV extracts were allowed to settle on poly-L lysine coated coverslips and then imaged on an Zeiss Imager Z2 with a 63X objective (N.A. 1.4) or with a SP5 confocal (Leica) with a 40X objective (N.A. 1.25).

### Spectroscopy

EVs labeled with either MemBright-Cy3 or PKH-26, or control MemBright-Cy3 or control PKH (diluted in PBS as described above), as well as the dyes directly diluted in Milli-Q water (Millipore) or ethanol were analysed by spectroscopy. Absorption and emission spectra were recorded at 20°C in quartz cuvettes on a Cary 400 Scan ultraviolet–visible spectrophotometer (Varian) and a FluoroMax-4 spectrofluorometer (Horiba Jobin Yvon) equipped with a thermostated cell compartment, respectively. For standard recording of fluorescence spectra, excitation was at 520 nm and the emission was collected 10 nm after the excitation wavelength (530 nm to 700 nm). All the spectra were corrected from wavelength-dependent response of the detector. The scattering due to the EVs was corrected with a baseline correction using Origin software. Quantum yields were determined using rhodamine B in water (QY= 0.31) as a reference (Magde et al., 1999).

### Fluorescence Correlation Spectroscopy (FCS)

To characterize the size of PKH aggregates, FCS measurements were performed on PKH26 (diluted at 5 μM) using a home-built confocal set-up based on a Nikon inverted microscope with a Nikon 60x 1.2NA water immersion objective. Excitation was provided by a cw laser diode (532 nm, Oxxius) and photons were detected with a fibered Avalanche Photodiode (APD SPCM-AQR-14-FC, Perkin Elmer) connected to an on-line hardware correlator (ALV7000-USB, ALV GmbH, Germany). Typical acquisition time was 5 min (10 × 30 s) with an excitation power of 1.1 μW at the sample level. The data were analyzed using the PyCorrFit software (Müller et al., 2014b).

### MemBright labeling of cells

Sub-confluent cells in 10cm culture dishes were rinsed twice with warm serum free medium and then incubated for 30 minutes at 28°C (Zmel1 cells) or at 37°C (4T1 cells) with MemBright quickly diluted in serum free medium (200nM final). To eliminate all possible traces of unbound MemBright, cells were then rinsed three times with serum free medium, rinsed with EDTA and trypsinated. Cells were then either injected in zebrafish embryos, seeded in triple flask for EV production, or seeded in glass bottom microwell dishes (MatTek Corporation) pre-coated with fibronectin from bovine plasma at 10μg/ml (Sigma F-1141) for imaging.

### Zebrafish handling

*Tg(fli1a:eGFP), Tg(mpeg1:eGFP)* and *Tg(mpo:eGFP)* Zebrafish (Danio rerio) embryos from a *Golden* background used in the experiments were kindly provided by F. Peri’s (EMBL, Heidelberg, Germany) and C. Lengerke’s laboratories (University Hospital Basel, Switzerland). Embryos were maintained at 28°C in Danieau 0.3X medium (17,4 mM NaCl, 0,2 mM KCl, 0,1 mM MgSO_4_, 0,2 mM Ca(NO_3_)_2_) buffered with HEPES 0,15 mM (pH = 7.6), supplemented with 200 μM of 1-Phenyl-2-thiourea (Sigma-Aldrich) to inhibit the melanogenesis, as previously described (Goetz et al., 2014).

### Intravascular injection of zebrafish embryo

At 48h post-fertilization (hpf), zebrafish embryos were dechorionated and mounted in 0.8% low melting point agarose pad containing 650 μM of tricain (ethyl-3-aminobenzoate-methanesulfonate) to immobilize them. Pre-labelled EVs, polystyrene beads or tumors cells were injected with a Nanoject microinjector 2 (Drummond) and microforged glass capillaries (25 to 30 μm inner diameter) filled with mineral oil (Sigma). 13,8 nL of a EV, beads or cell suspension (at 100.10^6^ cells) per ml were injected into the duct of Cuvier of the embryos under the M205 FA stereomicroscope (Leica), as previously described (Follain et al., 2018b; Stoletov et al., 2010). For late endosome/lysosome labeling, embryos were incubated with Lysotracker Deep Red (Thermo Fisher Scientific) diluted at 5μM in Danieau 0,3X medium for 2 hours at 28°C before injection.

### Confocal imaging and analysis

Confocal imaging was alternatively performed with an inverted TCS SP5 with HC PL APO 20X/0,7 IMM CORR CS objective (Leica) or an upright SP8 confocal microscope with a HC FLUOTAR L 25X/0,95 W VISIR objective (Leica). For high speed imaging of EVs in the blood flow, embryos were imaged right after injection; acquisitions were done at 80-100 frames per second for 1 minute, using the resonant scanner in a single Z plane, with an opened pinhole of more than 1 airy unit. To identify the cell types utapking EVs, the caudal plexus region of mpeg1:GFP, mpo:GFP or Fli1a:GFP was imaged 3h post-injection with a z-step of 1 μm . To quantify the proportion of EVs arrested in the dorsal aorta *vs* venous plexus regions, images were acquired similarly in Fli1:GFP embryos. For each case, quantification is described in the next paragraphs. To image the dynamics of macrophage protrusions, short time lapses of mpeg1:GFP embryos were acquired at 5 to 10 Z stacks per minute (z-step of 0,5 μm, stack covering the macrophage). To image the dynamics of macrophages, long time lapses of mpeg1:GFP embryos were acquired at 1 Z stack per minute for one hour in (z-step of 2 μm, stack covering the venous plexus). To image the uptake of EVs by macrophage, mpeg1:GFP embryos short time lapses were generated right after injection at 3 to 8 images per second on single Z planes. Image analysis and processing was performed using Fiji (Schindelin et al., 2012) as described in the following paragraphs.

### Semi-automated method to determine the proportion of internalized EVs

To determine the proportion of EVs internalized by either endothelial or macrophages, we used the Z-stacks obtained from either Fli1:GFP or mpeg1:GFP embryos injected with Zmel1-MemBright EVs. Using Fiji, we splitted the cell and EVs channels and merged them in a single RGB image. From the merged channel, we made a binary stack followed by a Z-projection with maximal intensity. We used this as a reference image where all the EVs and cells are apparent. After normalyzing this image to 1 we multiplied each stack (respectively EVs and Cell) by this projection. In both stacks we thus keep only the positions that colocalize either with the EV position or the Cells position (all other positions possess a null value). We then made a binary from the Cell stack, apply close and dilate before normalizing it to 1. The multiplication of this stack with the EV one lead to a new stack that keeps only the particle enclosed in the cellular compartments. Getting back to the Cell stack, we apply an inversion of the intensity values before substracting 254. The resulting stack is then multiplied by the EV stack and the created new stack let only appearent the EVs that are not colocalised with the cells. Further analyses of the intensities from the two stacks allowed us to access the ratiometric values of EVs uptaken by the different cell lines.

### Quantification of EVs in aorta Vs vein regions

Each region (dorsal aorta and venous plexus) was manually delimited on Z-projections, using vessels visible in Fli1:GFP channels. Total EV intensity was then measured in each region and reported to the area. A ratio of EV fluorescence in the venous plexus over dorsal aorta was then measured for each fish.

### Flow analysis for Red blood cells

#### Flow analysis of red blood cells

we first globally enhanced the contrast of the whole stack. Then we performed a Z-projection with the average intensity and substracted the obtained image to the stack. The remaining stack exhibits only the moving objects i.e. the red blood cells in this case. Then we applied a binarisation to the stack before applying a bandpass filter with the correct values to remove the background noise and keeping only the flowing blood cells. This stack is then further analysed with the Mosaic 2D/3D particle tracker plugin. We thus accessed the positions of each blood cell for the different frames and we computed the velocities of each individual track.

#### Flow analysis of EVs

Time-lapses of EVs were first tresholded and binarized. We then inverted the stack before running the 2D spot enhancing Filter plugin. We used the resulting stack to perform a second binarisation and then launched the Mosaic 2D/3D particle tracker plugin. We thus accessed the positions of each EV for the different frames and we computed the velocities of each individual track

### EVs and RBCs distance and velocity from the endothelial barrier

In order to access to the distance of the EVs or red blood cells to the endothelilal barrier, we first drew the endothelial wall using the transmitted light and extracted its coordinates to a table. From the analysis described in the previous paragraph, we extracted the coordinates and the velocity EVs and red blood cells. We ran a macro where we compared for all the position X_EV_ and Y_EV_ of the EV the closest postion X_endo_ and Y_endo_ by comparing all the possible distances d by calculating:

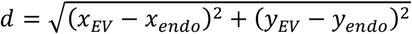

and keeping the smallest distance.

This allows us to plot the EV or the red blood cells velocities as a function of the distance from the endothelial wall.

### Sample preparation for Correlative Light and Electronic Microscopy of ZF embryos

Correlative Light and Electron Microscopy was performed as previously described (Goetz et al., 2014; Karreman et al., 2016a). Transgenic mpeg1:GFP embryos were injected with MemBright-Cy3 4T1 EVs and imaged alive with a Leica SP8 confocal microscope (see “Confocal imaging and analysis section”). Z stack was performed on two patrolling macrophages having uptaken EVs. After imaging, the embryo was chemically fixed with 2,5% glutaraldehyde and 4% paraformaldehyde in 0.1 M Cacodylate buffer (the fish tail was cut off in the fixative). Sample was kept in fixative at room temperature for 1-2h and stored in fixative at 4°C overnight or until further processing. Sample was rinsed in 0.1M Cacodylate buffer for 2×5min and post-fixed using 1% OsO4 in 0.1 M Cacodylate buffer, for 1h at 4°C. Then, sample was rinsed for 2×10min in 0.1M Cacodlyate buffer and secondary post-fixed with 4% water solution of uranyl acetate, 1h at room temperature. Rotation was used at all steps of sample processing. Followed by 5min wash in MiliQ water, the sample was stepwise dehydrated in Ethanol (25%, 50% each 15min, 95%, 3X100% each 20min) and infiltrated in a graded series of Epon (Ethanol/Epon 3/1, 1/1, 1/3, each 45min). Sample was left in absolute Epon (EmBed812) overnight. The following day, sample was placed in a fresh absolute Epon for 1h and polymerized (flat embedded) at 60°C for 24-48h. Once polymerized, most surrounding Epon was cut off using razorblade and sample was mounted on empty Epon blocks (samples flat on the top of the blocks) and left at 60 °C for 24h-48h. Samples were attached to an imaging pin with dental wax and mounted into the Brukker Skyscan 1272 for microCT imaging. Data was acquired over 188° with 0.2° angular step and a pixel size of 9 μm. Karreman et al. thoroughly details the process of how the microCT data enables the correlation of fluorescent imaging to 3D EM of voluminous samples (Karreman et al., 2016a). Retrieval of the region of interest in described in Supplementary Figure 8. The region of interest was targeted by ultramictrome, sections stained with toluidine blue and compared with the MicroCT and LM datasets. After targeting, serial 70nm sections were collected in formvar coated slot grids. The sections were post stained with uranyl acetate (4%) and lead citrate. The sections were imaged in a Biotwin CM120 Philips (FEI) TEM at 80kV with a SIS 1K KeenView. Stitches of the 70 sections were aligned using the Track EM pluggin in Fiji (Cardona et al., 2012). Segmentation and 3D reconstruction was done using the IMOD software package (Boulder Laboratory, University of Colorado) and Amira.

### Statistics

Statistical analysis of the results was performed using the GraphPad Prism program version 5.04. The Shapiro-Wilk normality test was used to confirm the normality of the data. The statistical difference of Gaussian data sets was analyzed using the Student unpaired two-tailed t test, with Welch’s correction in case of unequal variances. For data not following a Gaussian distribution, the Mann-Whitney test was used. Illustrations of these statistical analyses are displayed as the mean +/- standard deviation (SD). p-values smaller than 0.05 were considered as significant. *, p<0.05, **, p < 0.01, ***, p < 0.001, ****, p < 0.0001.

## Acknowledgments

We are indebted to Kerstin Richter and Francesca Peri (EMBL, Heidelberg, Germany) as well as to Pauline Hanns and Claudia Lengerke (University Hospital Basel, Switzerland) for supplying zebrafish embryos. We are grateful to R. White (MSKCC) for the Zmel1 (native and tdTomato) cells, as well as to Dr. Dorothy C. Bennett, (St. George’s University of London), and Dr. M. Soengas (CNIO, Madrid) for mammalian melanoma cells. We thank A. Michel (EFS) and C. Spiegelhalter (IGBMC) for electron microscopy image acquisition assistance, Elvire Guiot and Yves Lutz (IGBMC) for advices on confocal imaging and A. Audfray (Malvern Instruments) for NTA. We thank the CNIO proteomics core for performing the mass spectrometry on mouse and human melanoma models. We thank Philippe Herbomel for critical reading of the manuscript. This work was supported by a fellowship from IDEX (University of Strasbourg) to SG, by grants from La Ligue contre le Cancer, Canceropole Grand-Est and Roche to JG, and by institutional funds from University of Strasbourg, INSERM and ANR (French Proteomics Infrastructure ProFI; ANR-10-INBS-08-03).

## Author contributions

VH and JG planned the project. VH designed and conducted most of the experiments with contributions from SG and JB. MC synthesized the MemBright and led the spectroscopy experiments, with ASK. SH designed the automated EV tracking and the EV colocalization analysis. OL generated the 4T1 CD63-GFP line. FD, JB and CC conducted the mass spectrometry analysis on Zmel1 EVs. AIA, SGS and HP lead the mass spectrometry analysis on human and mice melanoma EVs. FV and GVN generated the mass spectrometry data on AB9 and endogenous zebrafish EVs. NF prepared the samples for the CLEM experiment and PM did the serial sectioning and the microCT. YS gave advices to conduct the CLEM experiment. LM and IB contributed to the segmentation and 3D modelling for the CLEM experiment. VH and JG wrote the manuscript with insights from all authors.

## Supplementary figure legends

**Supplementary Figure 1: Spectroscopy analysis of Zmel1 and 4T1 EVs labeled with MemBright or PKH. (A)** Histograms showing a spectroscopy analysis of MemBright and PKH describing the absorbance (left, y axis) and the fluorescence intensity (right, y axis) versus the wavelength (nm, x axis) of the two probes in water or methanol. The presence of aggregates of PKH in water is visible. Arrows indicate the presence of PKH aggregates in labeled EVs (left) as well as in control PKH alone (right). **(B)** Histograms showing the absorbance (left, y axis) and the normalized absorbance (right, y axis) of Zmel1 or 4T1 EVs labeled with PKH or MemBright versus the wavelength (nm, x axis). PKH aggregates are denoted with an arrow. **(C)** Histograms showing the intensity of the emitted fluorescence (left, y axis) and the normalized fluorescence intensity (right, y axis) of Zmel1 or 4T1 EVs labeled with PKH or MemBright versus the versus the wavelength (nm, x axis). PKH fluorescent aggregates are denoted with an arrow.

**Supplementary Figure 2: Microscopy comparison of MemBright and PKH labeled EVs. (A)** Representative fluorescent images of Zmel1 EVs labeled with PKH (at 2μM) or MemBright (at 200nM) and histogram showing the relative fluorescent intensity of individual puncta (p=0,001; Mann-Whitney test). **(B)** Representative fluorescent images of 4T1 EVs labeled with PKH (at 200nM) or MemBright (at 200nM) and histogram showing higher fluorescent intensity of Zmel1-MemBright individual puncta compared to Zmel1-PKH puncta (p<0,0001; Mann-Whitney test).

**Supplementary Figure 3: Density gradient of MemBright labeled 4T1 EVs.** Western blot on EVs labeled with MembrightCy3, or MemBright alone, separated on a density gradient (Left). It shows the presence of Alix and TSG-101 in the fractions 5-10 exclusively. No signal is observed in the control MemBright alone. Representative fluorescent images at low (upper) and high (lower) magnifications of the same samples than the westernblots (right). Fluorescent MemBrightCy3 puncta accumulate in fractions 5-10.

**Supplementary Figure 4: Size comparison of EVs and 100nm polystyrene beads. (A)** Representative confocal images of Zmel1 EVs labeled with MemBright-Cy5 and incubated with 100nm red fluorescent polystyrene beads *in vitro*. **(B)** Representative confocal Z projections of *Tg(pu1:GFP)* (lymphoid, monocytes/macrophages) embryos co-injected with Zmel1 EVs labeled with MemBright-Cy5 and with 100nm red fluorescent polystyrene beads imaged 3 hours post-injection. **(C)** Single plane zoom on embryos co-injected with Zmel1 EVs labeled with MemBright-Cy5 and with 100nm red fluorescent polystyrene beads. **(D)** Histogram showing the apparent diameters (left, nm) of MemBright labeled Zmel1 EVs and 100nm beads measured in confocal images *in vitro* and *in vivo* in zebrafish embryos *(in vitro:* p<0,0001; *in vivo:* p=0,6; Mann-Whitney test).

**Supplementary Figure 5: Dual color EV MemBright labeling. (A)** Representative confocal Z projections of *Tg(Fli1:GFP)* embryos co-injected with Zmel1 EVs labeled with MemBright-Cy3 and with 4T1 EVs labeled with MemBright-Cy5. **(B)** Representative confocal single planes from a time-lapse imaged right after injection of *Tg(Fli1:GFP)* embryos co-injected with Zmel1 EVs labeled with MemBright-Cy3 and with 4T1 EVs labeled with MemBright-Cy5. (C) Time projection over 10 seconds of a time-lapse imaged right after injection of *Tg(Fli1:GFP)* embryos co-injected with Zmel1 EVs labeled with MemBright-Cy3 and with 4T1 EVs labeled with MemBright-Cy5.

**Supplementary Figure 6: Control Zebrafish embryo injected with MemBright-labeled EVs or with control MemBright alone.** Representative confocal Z-projections of *Tg(mpeg1:GFP)* (macrophages) embryos injected with either 4T1 EVs labeled with MemBright-Cy3 or with MemBright-Cy3 without EVs and imaged 3 hours post injection.

**Supplementary Figure 7: Velocity of patrolling macrophages. (A)** Histogram showing the velocity of non-injected *Tg(mpeg1:GFP)* positive macrophages (y axis, μm/min) in relation with their perimeter (x axis, μm). **(B)** Left: histogram showing the intensity of uptaken EVs (y axis, A.U.) as a function of the macrophages velocity (x axis, μm/min) in *Tg(mpeg1:GFP)* embryos injected with MemBright labelled 4T1 EVs. Middle: histogram showing the average intensity of uptaken EVs (y axis, A.U.) depending on the macrophage perimeter (x axis, μm) (p=0,002; Student t-test). Right: histogram showing the average macrophages velocity (y axis, μm/min) depending on their perimeter (x axis, μm) (p=0,005; Mann-Whitney). **(C)** Left: histogram showing the intensity of uptaken EVs (y axis, A.U.) as a function of the macrophages velocity (x axis, μm/min) in *Tg(mpeg1:GFP)* embryos injected with MemBright-labelled Zmel1 EVs. Right: histogram showing the macrophage perimeter (y axis, μm) as a function of the macrophages velocity (x axis, μm/min) in *Tg(mpeg1:GFP)* embryos injected with MemBright-labelled Zmel1 EVs.

**Supplementary Figure 8: Retrieval of the cells by CLEM. (A)** *Tg(mpeg1:GFP)embryos* were injected with 4T1 MemBright-Cy3 labeled EVs and imaged by confocal (upper panels). The upper right panel shows the position of the Region Of Interest (ROI) containing the two target cells, with respect to several embryonic landmarks imaged by confocal at low magnification. The lower left image shows the tail of the embryo after fixation and resin embedding imaged by microCT. The lower right image shows the position of the ROI in an electron microscopy section. **(B)** Higher magnification of the ROI imaged by confocal and electron microscopy. Common features between transmitted light in the living fish and electron microscopy on fixed fish are highlighted to allow a precise positioning of the ROI. The asterisk points to a dirt present on the EM section.

**Supplementary Figure 9: The putative journey of EVs in macrophages by electron microscopy. (A)** Electron microscopy images of EVs observed in the lumen of the vessel, in the close proximity of protrusions extending from the macrophage plasma membrane, which were identified by CLEM. **(B)** Electron microscopy images of putative EVs present in early endosomes close to the surface of macrophages. **(C)** Electron microscopy images of putative EVs present in MVBs.

**Supplementary Figure 10: 4T1 CD63-GFP cells pre-labelled with MemBright. (A)** Representative confocal images of 4T1 CD63-GFP cells labeled with MemBright-Cy3 at different times before and after MemBright addition. **(B)** Zooms on confocal images of 4T1 CD63-GFP cells labeled with MemBright-Cy3 at 3h and 24h after MemBright addition. **(C)** Representative images of EVs isolated from the extracellular medium of 4T1 CD63-GFP cells pre-labeled with MemBright-Cy3.

**Supplementary tables**

**Table 1:** proteins identified in Zmel1 EVs by mass spectrometry

**Table 2:** proteins identified in human melanoma (451-LU, SK-Mel28, SK-Mel147, SK-Mel103, WM35, WM164) EVs by mass spectrometry

**Table 3:** proteins identified in mouse melanoma EVs (B16-F0, B16-F1, B16-F10) by mass spectrometry

**Table 4:** proteins common to zebrafish, mouse and human melanoma and melanocyte EVs

**Table 5:** proteins common to Zmel1 EVs and AB9 EVs

**Table 6:** proteins common to Zmel1 EVs and YSL CD63-GFP positive EVs

**Table 7:** Quantum yield of MemBright and PKH labeled EVs

**Supplementary Movies**

**Movie 1 (related to Figure 3):** MemBright-labeled Zmel1 EVs imaged in the circulation of the caudal plexus right after injection.

**Movie 2 (related to Figure 3):** MemBright-labeled 4T1 EVs imaged at high speed in the dorsal aorta right after injection.

**Movie 3 (related to Figure 3):** MemBright-labeled 4T1 EVs imaged at high speed in the caudal vein right after injection.

**Movie 4 (related to Figure 4):** MemBright-labeled Zmel1 EVs imaged in the caudal vein right after injection in *Tg (Fli1:GFP)* embryos. Endothelium is green, EVs are red.

**Movie 5 (related to Figure 5):** *Tg(mpeg1:GFP)* positive macrophages present in the circulation of the caudal plexus and scanning the blood flow with their protrusions.

**Movie 6 (related to Figure 5 and Figure S7):** Dynamics of small round intravascular macrophages and more elongated ones in the caudal plexus of *Tg(mpeg1:GFP)embryos*.

**Movie 7 (related to Figure 5, 6 and Figure S8, S9):** CLEM experiment. Serial sections of 2 patrolling macrophages having uptaken EVs imaged by transmitted electron microscopy. 3D segmentation model of the macrophages (green), the endothelium (purple), circulating red blood cells (blue) and late endosomes / lysosomes within the macrophages (orange and red).

**Movie 8 (related to Figure 6):** MemBright-labeled Zmel1 EVs injected in *Tg(mpeg1:GFP)embryos* and imaged right after injection in the caudal plexus (large field view, Z projection). Macrophages are green, EVs are cyan.

**Movie 9 (related to Figure 6):** EV uptake by endocytosis. MemBright-labeled Zmel1 EVs injected in *Tg(mpeg1:GFP)* embryos and imaged right after injection in the caudal plexus (zoom, single plane). Macrophages are green, EVs are red.

**Movie 10 (related to Figure 6):** EV uptake by filopodia surfing / macropinocytosis. MemBright-labeled Zmel1 EVs injected in *Tg(mpeg1:GFP)* embryos and imaged right after injection in the caudal plexus (zoom, single plane). Macrophages are green, EVs are red.

**Movie 11 (related to Figure 7):** EVs from Zmel1 cells pre-labelled with MemBright are transferred to macrophages *in vivo. Tg(mpeg1:GFP)* embryos are injected in the circulation with tdTomato – Zmel1 cells pre-labelled with MemBright-Cy5 and imaged 48h post-injection in the caudal plexus. Macrophages are green, Zmel1 cells are red and MemBright is cyan.

